# Chemical-genetic profiling reveals cross-resistance and collateral sensitivity between antimicrobial peptides

**DOI:** 10.1101/542548

**Authors:** Bálint Kintses, Pramod K. Jangir, Gergely Fekete, Mónika Számel, Orsolya Méhi, Réka Spohn, Lejla Daruka, Ana Martins, Ali Hosseinnia, Alla Gagarinova, Sunyoung Kim, Sadhna Phanse, Bálint Csörgo, Ádám Györkei, Eszter Ari, Viktória Lázár, Anikó Faragó, László Bodai, István Nagy, Mohan Babu, Csaba Pál, Balázs Papp

**Author notes:** These authors contributed equally to this work (Bálint Kintses, Pramod K. Jangir, Gergely Fekete).

## Abstract

Antimicrobial peptides (AMPs) are key effectors of the innate immune system and promising therapeutic agents. Yet, knowledge on how to design AMPs with minimal cross-resistance to human host-defense peptides remains limited. Here, with a chemical-genetic approach, we systematically assessed the resistance determinants of *Escherichia coli* against 15 different AMPs. Although generalizations about AMP resistance are common in the literature, we found that AMPs with different physicochemical properties and cellular targets vary considerably in their resistance determinants. As a consequence, collateral sensitivity effects were common: numerous genes decreased susceptibility to one AMP while simultaneously sensitized to others. Finally, the chemical-genetic map predicted the cross-resistance spectrum of laboratory-evolved human-B-defensin-3 resistant lineages. Our work substantially broadens the scope of known resistance-modulating genes and explores the pleiotropic effects of AMP resistance. In the future, the chemical-genetic map could inform efforts to minimize cross-resistance between therapeutic and human host AMPs.

## Introduction

Antimicrobial peptides (AMPs) play a crucial role in general defense mechanisms against microbial pathogens in all classes of life. Although there is a considerable diversity in their amino acid content, length, and structure, AMPs are typically positively charged and amphipathic molecules^1,2^. These properties allow them to adsorb onto the bacterial cell surface and penetrate through the membrane to exert their diverse antibacterial actions^3^. As AMPs have a broad spectrum of activity, considerable efforts have been allocated to the research and development of novel anti-infective compounds originating from AMPs^4,5^. However, the clinical development of AMP therapies, has also raised concerns that these approaches may drive bacterial evolution of resistance to human host-defense peptides^6,7^. As well, therapeutic AMPs are required to be active against pathogenic bacteria, many of which have already evolved resistance against human host AMPs^8^. Therefore, ideally, resistance mechanisms against therapeutic and host AMPs should not overlap.

Accumulating evidence suggest that AMPs differ considerably in their mode of actions, which may influence the specific microbial resistance mechanisms against them^1,9^. First, there are substantial differences in the electrostatic interactions and transport processes that lead to the cellular uptake of AMPs^3,10^. Second, the cellular targets of AMPs are also diverse in nature. For instance, apart from their membrane-disruptive activities, AMPs inhibit intracellular processes such as bacterial DNA and RNA synthesis, translation, cell wall synthesis, and diverse metabolic pathways^1,11^. However, the extent to which the genetic determinants of resistance differ across AMPs remains unclear, because most of our knowledge comes from case studies characterizing only a limited number of membrane-targeting AMPs^9^ (Supplementary Table 1). Therefore, there is an urgent need to comprehensively map the relationships between the modes of action of AMPs and the genetic determinants influencing bacterial susceptibility to them. Understanding these complex relationships would help to rationally choose AMPs for clinical development which are dissimilar to human host peptides in terms of the underlying resistance mechanisms.

**Table 1.** List of AMPs used in this study, their abbreviation, described mode of action, and clinical relevance (for details see Supplementary Table 10).

| <u>Name of AMP</u> | <u>Abbreviation</u> | <u>Mode of action</u> | <u>Clinical relevance</u> |
| --- | --- | --- | --- |
| Apidaecin IB | AP | Inhibits protein biosynthesis by targeting ribosomes;<br>Interacts with DnaK, GroEL/GroES, FtsH | Yes |
| Bactenecin 5 | BAC5 | Inhibits protein and RNA synthesis | n.a. |
| CAP18 | CAP18 | Disrupts cell membrane | Yes |
| Cecropin P1 | CP1 | Disrupts cell membrane | n.a. |
| Human beta-defensin-3 | HBD-3 | Disrupts cell membrane;<br>Inhibits lipid II in peptidoglycan biosynthesis | n.a. |
| Indolicidin | IND | Inhibits DNA and protein synthesis;<br>Disrupts cell membrane;<br>Inhibits septum formation | Yes |
| LL-37 human cathelicidin | LL37 | Disrupts cell membrane;<br>Induces ROS formation | Yes |
| Peptide glycine-leucine amide | PGLA | Disrupts cell membrane | n.a. |
| Pexiganan | PEX | Disrupts cell membrane | Yes |
| Pleurocidin | PLEU | Disrupts cell membrane;<br>Induces ROS formation;<br>Inhibits protein and DNA synthesis | n.a. |
| Polymyxin B | PXB | Disrupts cell membrane;<br>Induces ROS formation | Yes |
| PR-39 | PR39 | Inhibits protein and DNA synthesis | n.a. |
| Protamine | PROA | Affects cellular respiration and glycolysis;<br>Disrupts cell envelop | n.a |
| R8 | R8 | n.a | n.a |
| Tachyplesin II | TPII | Disrupts cell membrane | n.a |

n.a. – no data available

Chemical-genetic profiling is a reverse genetic approach that quantifies the susceptibility of a genome-wide collection of mutant libraries to a set of chemical compounds^12^. By modulating gene dosage (i.e. either by depletion or overexpression), several studies demonstrated the effectiveness of this tool to map cellular targets and genetic determinants of resistance for antibiotics^13–18^. Moreover, antibiotics with similar chemical-genetic profiles, i.e. those with a large overlap between the gene sets influencing resistance to them, are likely to share cellular targets and mechanism of action^16^. Therefore, chemical-genetics has been proposed as a useful tool to infer if resistance evolution to an antibiotic would lead to cross-resistance (decreased sensitivity) or collateral sensitivity (increased sensitivity) to another antibiotic^12,19^.

Here we employed a genome-wide chemical-genetic approach to explore the diversity of resistance determinants across AMPs in the model bacterium *Escherichia coli* (*E. coli*). First, we generated a comprehensive chemical-genetic map by measuring how overexpressing each of ∼4,400 genes of *E. coli* influences the bacterium’s susceptibility against 15 AMPs. The set of 15 AMPs are structurally and chemically diverse and include AMPs with well-characterized modes of action, clinical relevance, or crucial role in the human immune defense (Table 1). By analyzing the large number of genes that influenced bacterial susceptibility to AMPs in the chemical-genetic screen, we identified major differences in the genetic determinants of resistance that clustered the AMPs according to their modes of action. Next, we confirmed our results with a complementary chemical-genetic approach by testing the growth effect of a smaller set of 4 selected AMPs against an array of 279 partially-depleted essential genes (i.e. hypomorphs)^20–22^. The latter approach provides information on the intrinsic resistome, i.e. the collection of genes that contribute to resistance at their native expression levels. Together, these screens revealed numerous genes that modulate susceptibility against membrane-targeting and intracellular-targeting AMPs in an antagonistic manner. Finally, evolving *E. coli* in the laboratory to become resistant to a key human host-defense AMP demonstrated that chemical-genetic profiles are predictive of cross-resistance patterns between AMPs.

## Results

### Chemical-genetic profiling reveals AMP resistance-modulating gene sets

We generated chemical-genetic interaction profiles for a diverse set of AMPs (Table 1) by screening them against a comprehensive library of gene overexpressions in *E. coli*^23^. Increasing gene dosage is a widely applied approach to reveal the targets of small molecule antibiotics^24,25^. It also informs on the ‘latent resistome’, that is, the collection of genes where a change from native expression level enhances resistance to a particular drug^26^. We applied a sensitive competition assay by monitoring growth of a pooled plasmid library overexpressing all the *E. coli* ORFs (Figure 1a), as we reported earlier^27^. Specifically, *E. coli* cells carrying the pooled plasmid collection were grown in the presence or absence of one of the 15 AMPs tested, at a sub-inhibitory concentration that increased the doubling time of the whole population by 2-fold. Following 12 generations of growth, the plasmid pool was isolated from each selection and the relative abundance of each plasmid was determined by a deep sequencing readout (see Methods). By comparing plasmid abundances in the presence and absence of each AMP, we calculated a chemical-genetic interaction score (fold-change value) for each gene and identified genes that increase or decrease susceptibility upon overexpression (Figure 1a, Supplementary Table 2, see Methods).

**Figure 1.**
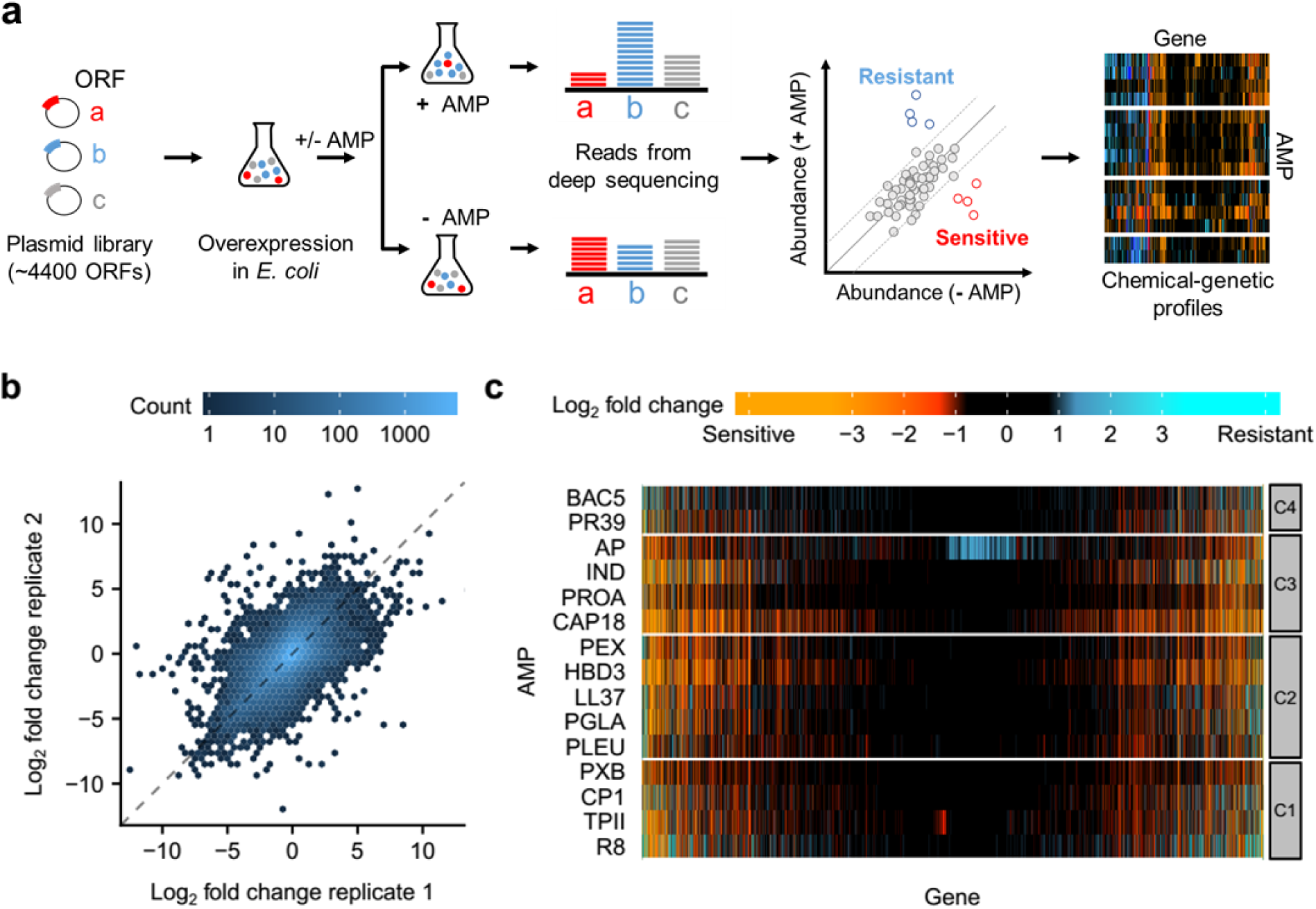
Chemical-genetic profiling of AMPs. a, Schematic representation of the chemical-genetic pipeline. The chemical-genetic interactions of ∼4,400 single gene-overexpressions and 15 different AMPs were measured using a pooled fitness assay with a deep sequencing readout (see Methods). **b,** A density scatter plot showing the overall correlation of replicate measurements of the chemical-genetic scores (log_2_ fold-change in the relative abundance of each gene in the presence vs absence of each AMP) across all genes and AMPs (r = 0.63 and *P* = 2.2*10^-16^, Pearson’s correlation, *n* = 53,292). **c,** Heatmap showing the chemical-genetic interaction scores. Resistant and sensitive interactions are represented by blue and red, respectively (*n* = 66,615). Groups C1-C4 refer to clusters defined in Figure 2. Data is provided in Supplementary Table 2.

To validate our workflow, we took three distinct approaches. First, we tested the reproducibility of the chemical-genetic profiles by correlating the chemical-genetic interaction scores between replicate measurements. The overall correlation was comparable to what has been achieved with arrayed mutants on high-density agar plates^16,28^ (r = 0.63 from Pearson’s correlation, Figure 1b). This indicates that we measured the growth effects with sufficiently high confidence. Second, we picked 15 overexpression plasmids that showed diverse chemical-genetic interaction scores with multiple AMPs in our screen but did not influence the growth rate of *E. coli* in the absence of AMPs (see Methods). Performing minimum inhibitory concentration (MIC) measurements confirmed 84% of these chemical-genetic interactions (Supplementary Figure 1 and Supplementary Table 3). Third, we collected examples from the literature where overexpression of an *E. coli* gene has been shown to influence sensitivity to a specific AMP. Despite differences in the used strains and protocols, 69% (9 out of 13) of the literature-curated interactions were captured by our screen (Supplementary Table 4). Taken together, these analyses indicate that our workflow has high sensitivity and is suitable to measure chemical-genetic interactions between AMPs and gene overexpressions.

### Chemical-genetic profiles group AMPs with similar mechanistic and physicochemical features

We first explored how similarity in the chemical-genetic profiles informs on the functional and physicochemical properties of AMPs. To this end, we compiled literature data on known modes of action (Table 1) and computed physicochemical properties for each AMP (see Methods and Supplementary Table 5). Next, we grouped peptides with similar chemical-genetic profiles using a robust clustering method (see Methods). This procedure resulted in four main clusters, referred to as C1 – C4 (Figures 1c and 2a).

We found that clusters C1 and C2 contain mostly AMPs that target primarily the bacterial membranes, whereas most AMPs in clusters C3 and C4 have intracellular targets (Figure 2a and Table 1). Membrane-targeting AMPs (C1 and C2) have unique physicochemical properties (Supplementary Figure 2). Specifically, they have a lower isoelectric point and proline content, they are substantially more hydrophobic and have a higher propensity to form secondary structures than C3 and C4 peptides (Figure 2b). These properties facilitate efficient integration of these AMPs into the bacterial membrane where they create pores^29,30^. Notably, although peptides in both C1 and C2 are pore-formers, they indeed show subtle differences in their physicochemical features when multiple properties are considered jointly (Supplementary Figure 3).

**Figure 2.**
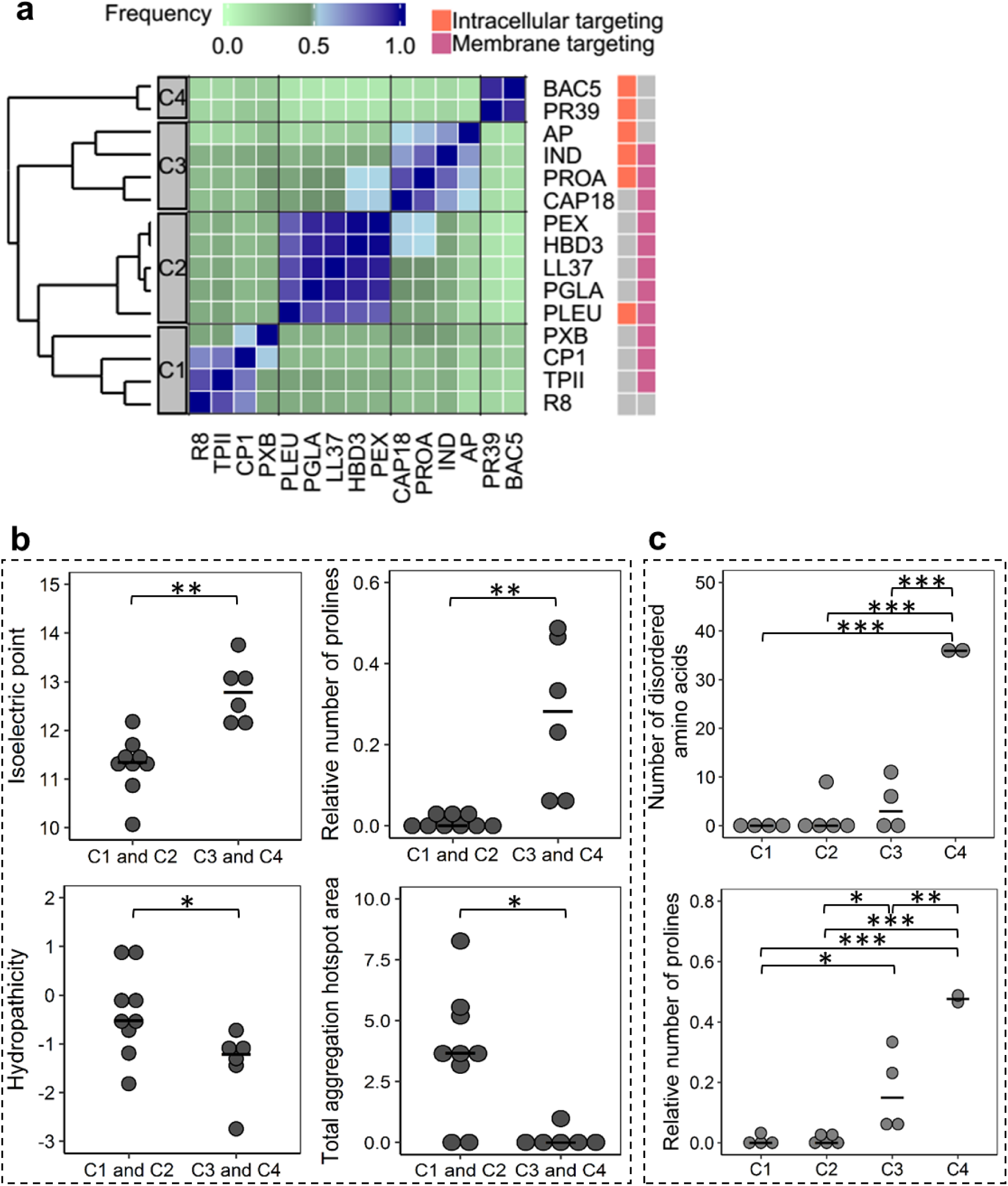
Chemical-genetic profiling discriminates membrane-targeting and intracellular-targeting AMPs with distinct physicochemical properties. **a,** Heatmap showing the ensemble clustering of the AMPs based on their chemical-genetic profiles (see Methods). For each AMP pair, the color code represents the frequency of being closest neighbours across the ensemble of clusters (*n* = 75,000 clustering). The four major clusters are labelled as C1, C2, C3, and C4. Membrane-targeting and intracellular-targeting broad modes of action are labelled with pink and orange, respectively, on the rightmost side of the figure. Grey color indicates that the specific broad mode of action has not been detected or not tested (see Table 1). References describing these activities are provided in Supplementary Table 10. **b,** Most important physicochemical properties that differentiated AMPs in cluster C1,C2 from AMPs in cluster C3,C4. Significant differences: ** *P* = 0.0026 and 0.0012 for isoelectric point and relative number of prolines, respectively, * *P* = 0.0391 and *P* = 0.0154 for hydropathicity and total aggregation hotspot area, respectively, two-sided Mann–Whitney U test, *n* = 9 and *n* = 6 for C1,C2 and C3,C4, respectively. **c,** Physicochemical properties that distinguished the clusters when the 4 main AMP clusters were considered separately (p<0.05 ANOVA, Tukey post-hoc test, *n* = 15). Significant differences: *** *P* = 1.1*10^-6^, *P* = 1.3*10^-6^ and *P* = 4*10^-6^ for C1 vs C4, C2 vs C4 and C3 vs C4, respectively in the case of number of disordered amino acids. * *P* = 0.034 and *P* = 0.027 for C1 vs C3 and C2 vs C3, respectively. ** *P* = 0.0022 for C3 vs C4. *** *P* = 5.5*10^-5^ and *P* = 4.2*10^-5^ for C1 vs C4 and C2 vs C4, respectively, in the case of relative number of prolines. Central horizontal bars represent median values. Data is provided in Supplementary Table 5.

The two clusters of intracellular-targeting AMPs (C3 and C4) have distinct physicochemical properties. In particular, AMPs in cluster C4 have an especially high proline content, leading to elevated propensity to intrinsic structural disorder (Figure 2c). Structural disorder has been described as a common feature in a novel class of intracellular-targeting AMPs^31^. Indeed, the two AMPs in cluster C4 Bactenecin 5 (BAC5) and cathelicidin PR-39 are known to have intracellular targets only as they do not lyse the membrane (Table 1). By contrast, AMPs in cluster C3 showed features of both membrane- and intracellular-targeting ones (Figure 2). Reassuringly, Indolicidin (IND) and Protamine (PROA), which are in cluster C3, have been described to have both membrane disruptive and intracellular-targeting activities (Table 1). Finally, while CAP18 is generally considered as membrane-targeting, our data indicate that it could also have intracellular targets as it clusters with PROA in the chemical-genetic map (Figure 2a). Future works should elucidate the exact mode of action of this peptide.

Taken together, AMPs with similar chemical-genetic profiles share physicochemical features and previously described broad mechanisms of action, indicating that chemical-genetics can capture certain differences in the bactericidal effects across AMPs.

### Large and functionally diverse set of genes influences AMP resistance

We identified between 88 – 778 and 348 – 1263 genes enhancing resistance and sensitivity per AMP, respectively (Figure 3a). This finding substantially broadens the scope of genes that enhance resistance (latent resistome), as previously reported resistance genes in *E. coli* constitute only 0.46% (11 of 2371) of the resistance-conferring genes identified here (Supplementary Table 6). Importantly, whereas genes annotated with cell envelope function were overrepresented among AMP susceptibility modulating genes (Supplementary Table 7), the majority of our hits did not have obvious functional connection with known AMP uptake mechanism or mode of action (Supplementary Figure 4).

**Figure 3.**
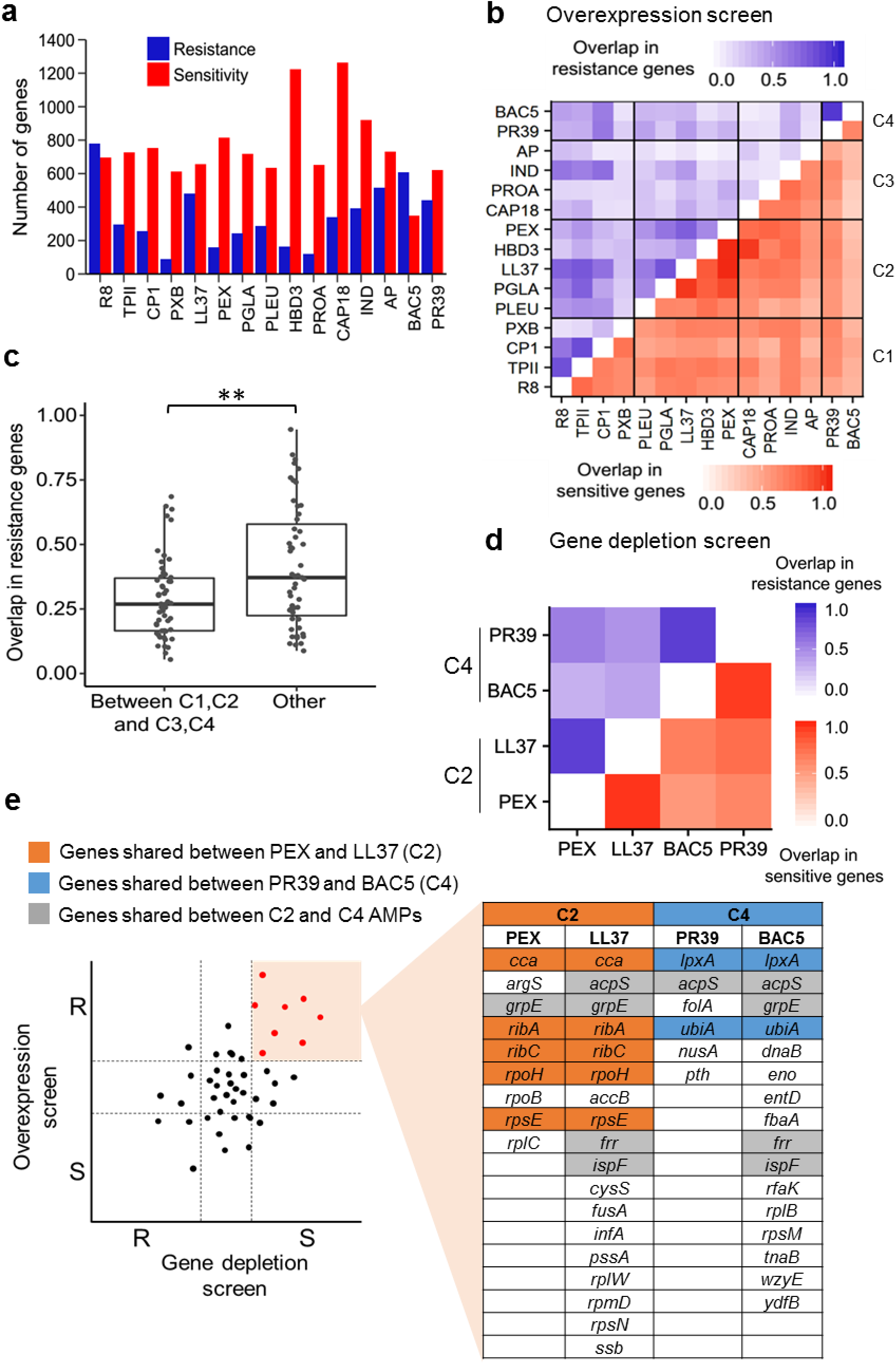
Large and functionally diverse latent and intrinsic AMP resistomes. **a,** Number of genes enhancing resistance and sensitivity for each AMP upon overexpression. Data is provided in Supplementary Table 2. **b,** Heatmap shows the corrected Jaccard similarity indices calculated for resistance(blue) and sensitivity-conferring genes (red) between AMP pairs based on the overexpression screen (see Methods for calculation of corrected Jaccard indices, *n* = 210, that is, the number of AMP pairs used for calculating the Jaccard-similarity indices). The darker the color the higher the overlap of gene sets between AMP pairs. Data is provided in Supplementary Table 6. **c,** The overlaps in the latent resistomes (genes conferring resistance upon overexpression) between AMP pairs belonging to different chemical-genetic clusters. Significant difference: ** *P* = 0.009 from two-sided Mann–Whitney U test, *n* = 54 and *n* = 51 for between C1, C2 and C3, C4, and others, respectively. **d,** Heatmap shows the corrected Jaccard similarity indices calculated for resistant (blue) and sensitive (red) chemical-genetic interactions with partially-depleted essential genes (see Methods). Data is provided in Supplementary Table 8. **e,** Sets of essential genes that simultaneously confer AMP resistance when overexpressed and sensitivity when depleted (red colored dots in the schematic plot). Color code is explained in the figure.

Next, to assess the diversity of resistance determinants across AMPs, we calculated the extent to which the resistance- and sensitivity-conferring genes are shared between pairs of AMPs. To avoid underestimating the overlap between gene sets across AMPs, we employed an index of overlap that takes into account measurement noise (see Methods). Typically, ∼63 % of the sensitive and ∼31 % of the resistance genes overlapped between pairs of AMPs (Supplementary Figure 5). The latter figure indicates substantial variation in the latent resistome across AMPs. Remarkably, the sets of resistance-conferring genes varied greatly even between AMPs in the same chemical-genetic cluster, in particular between AMPs in cluster C3 (Figure 3b). This pattern could reflect subtle differences in the modes of action across the intracellular-targeting AMPs within cluster C3 as these peptides differ in their specific targets (Table 1). Indeed, on a broader scale, membrane-targeting AMP pairs (C1-C2) and intracellular-targeting AMP pairs (C3-C4) shared more resistance genes than AMP pairs with different broad mechanisms of action (Figure 3c).

These findings reveal a vast diversity of resistance determinants across peptides that reflects differences in their modes of action and specific targets.

### Partial depletion of essential genes reveals intrinsic resistance to AMPs

Chemical-genetic profiling based on gene depletion captures a different aspect of resistance determinants than gene overexpression^32^. While resistance upon increased gene dosage informs on the latent resistome, hypersensitivity upon gene depletion reveals genes that contribute to resistance at their native expression levels, collectively called as the intrinsic resistome^26^. To investigate the intrinsic AMP resistome, we initiated a chemical-genetic screen with a set of 279 partially-depleted essential genes (hypomorphic alleles) of *E. coli*. We selected four AMPs with well-characterized modes of action, including two membrane-targeting (Pexiganan (PEX) and LL37 from C2) and two intracellular-targeting AMPs (BAC5 and PR39 from C4). Then, using a well-established high-density agar plate assay^21,22,33^, we determined their chemical-genetic interaction profiles across the hypomorphic alleles (Methods, Supplementary Table 8).

In total, we found that 75% of the 279 partially-depleted essential genes influenced susceptibility to at least one of the AMPs studied and 60% of these interactions caused hypersensitivity, indicating that essential genes often contribute to the intrinsic AMP resistome (Supplementary Table 8). We found substantial overlaps in the intrinsic resistomes between AMPs with similar modes of action. As high as 87% of the 279 hypomorphic alleles overlapped between PEX and LL37, and a similar figure emerged from the comparison of the gene set between BAC5 and PR39 (Figure 3d). In contrast, on average, only 59% of the 279 hypomorphic alleles were identical when functionally dissimilar AMPs were compared (Figure 3d).

Genes that simultaneously confer drug resistance when overexpressed and sensitivity when depleted are of special interest as such genes are likely to directly protect bacteria against drug stress or encode drug targets^34^. Comparison of our overexpression and hypomorphic screens revealed dozens of essential genes that showed both properties (Figure 3e). Remarkably, *folA* (dihydrofolate reductase), a known intracellular target of PR39^35^, was among the set of 6 genes that simultaneously conferred resistance when overexpressed and sensitivity when depleted in the presence of PR39. Similarly, *pssA* (a phosphatidylserine synthase) appeared in the presence of LL37, a membrane-targeting AMP. Reassuringly, deletion of *pssA* has been shown to alter membrane properties and increase bacterial sensitivity to membrane-targeting peptides^36^.

Together, these results indicate that both the intrinsic and the latent AMP resistomes are vast and shaped by the AMP’s mode of action.

### Collateral sensitivity interactions are frequent between AMPs with different modes of action

The limited overlap in resistance determinants across AMPs prompted us to hypothesize that some of the gene overexpressions might even have antagonistic effects against distinct AMPs. Specifically, we sought to identify resistance genes that induce collateral sensitivity, i.e. increase resistance to one AMP while simultaneously sensitize to another one^37,38^. We found numerous such cases (Supplementary Table 6). For example, out of the 4,400 genes, we retrieved 643 that conferred resistance to 2 or more AMPs while increasing sensitivity to at least 2 other AMPs upon overexpression.

For each pair of AMP, we then calculated the overrepresentation of collateral sensitivity-inducing genes over random expectation (see Methods). Intriguingly, pairs of AMPs within the same chemical-genetic cluster were typically depleted in such genes (Figure 4a). In contrast, the relative overrepresentation of collateral sensitivity-inducing genes was pronounced between the clusters of membrane-targeting and intracellular-targeting AMPs (Figure 4b). Finally, both the overexpression and the hypomorphic allele screens indicate that collateral sensitivity interactions are prevalent between proline-rich AMPs in cluster C4 (BAC5, PR39) and membrane-targeting AMPs (Figure 4a and Supplementary Figure 6).

**Figure 4.**
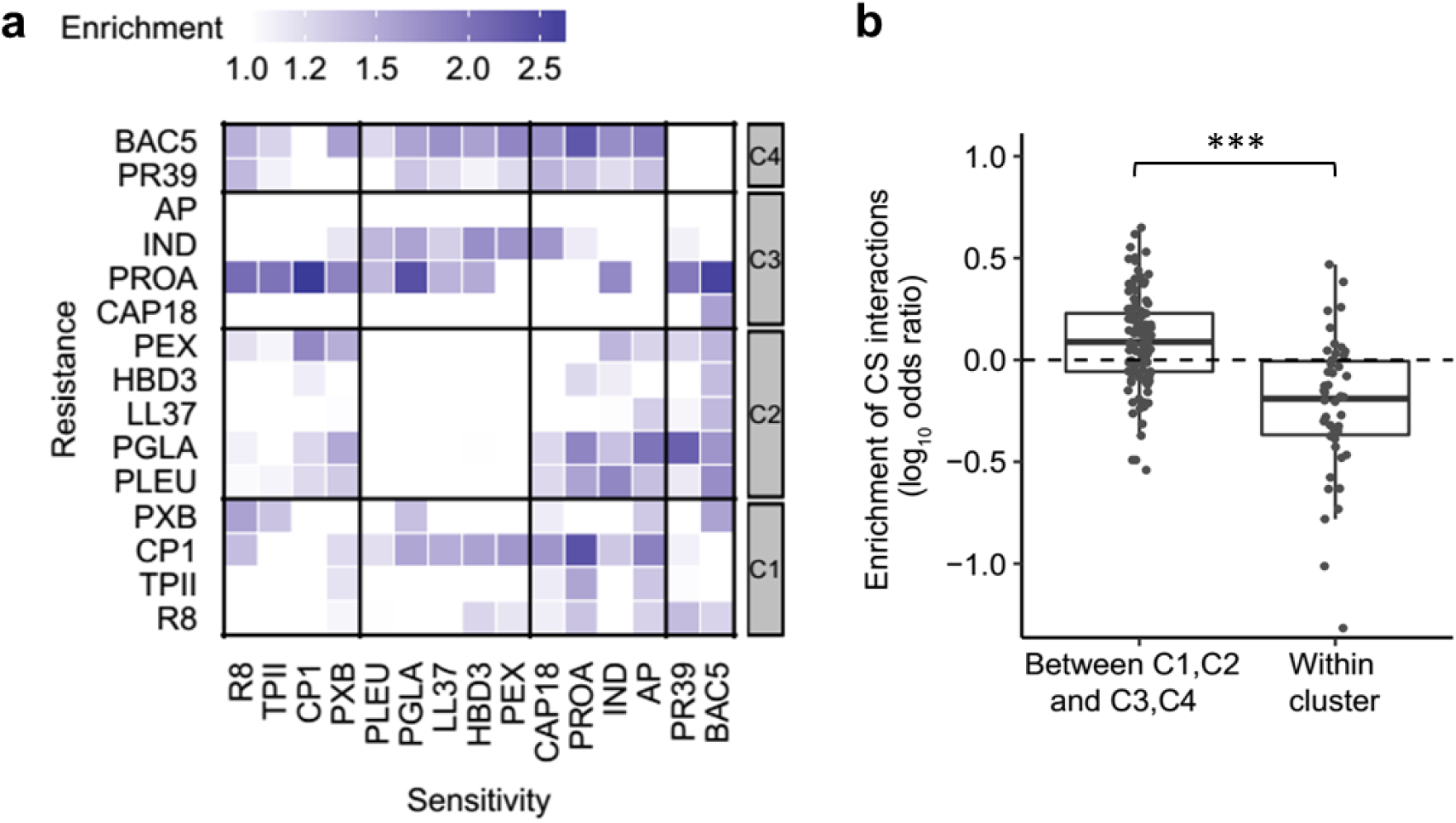
Collateral sensitivity (CS) interactions are frequent between AMPs with different modes of action. **a,** Heatmap depicting the overrepresentation of collateral sensitivity-inducing genes for each AMP pair over random expectation (*n* = 210 AMP pairs). Random expectation is calculated using the number of resistance and sensitive genes for each AMP (see Methods). **b,** Collateral sensitivity effects were especially pronounced between AMP pairs with different broad mechanism of action, that is, between membrane-targeting (C1, C2) and intracellular-targeting (C3, C4), as compared to AMP pairs from the same cluster. Significant difference: *** *P* = 2.1*10^07^ from two-tailed unpaired *t*-test, *n* = 108 and 46 for pairs of AMPs between C1, C2 and C3, C4, and those within cluster, respectively. Y-axis shows odds ratio (log10) of enrichment of collateral sensitivity interactions between AMP pairs. Data is provided in Supplementary Table 6.

### Perturbed phospholipid trafficking as a mechanism of collateral sensitivity

We next focused on genes that showed reduced susceptibility to at least 4 membrane-targeting AMPs upon overexpression while at the same time showed elevated susceptibilities toward at least 4 intracellular-targeting AMPs. These genes were enriched in functions related to phospholipid and lipopolysaccharide (LPS) composition of the bacterial membrane (Supplementary Figure 7). This trend is exemplified by MlaD and MlaE proteins (Supplementary Figure 7a), both being part of a protein complex that carries out retrograde phospholipid transport from the outer membrane to the inner membrane in Gram-negative bacteria^39^. Importantly, several studies have reported a role of the Mla (maintenance of lipid asymmetry) pathway in bacterial pathogenesis, virulence and antibiotic resistance^40–42^.

What could be the mechanism behind the antagonistic action of this pathway on membrane- versus intracellular-targeting AMPs? Since MlaD is part of a protein complex, it may lead to a loss-of-function effect upon overexpression^43,44^. To test this, we asked whether overexpression and deletion of *mlaD* cause similar changes in susceptibility to a representative set of membrane- and intracellular-targeting AMPs. Both mutations caused a decreased susceptibility to membrane-targeting AMPs and an increased susceptibility to intracellular-targeting ones (Figure 5a, for MIC curves, see Supplementary Figure 8 and 9), demonstrating that overexpression perturbs *mlaD* function similar to a loss-of-function mutation.

**Figure 5.**
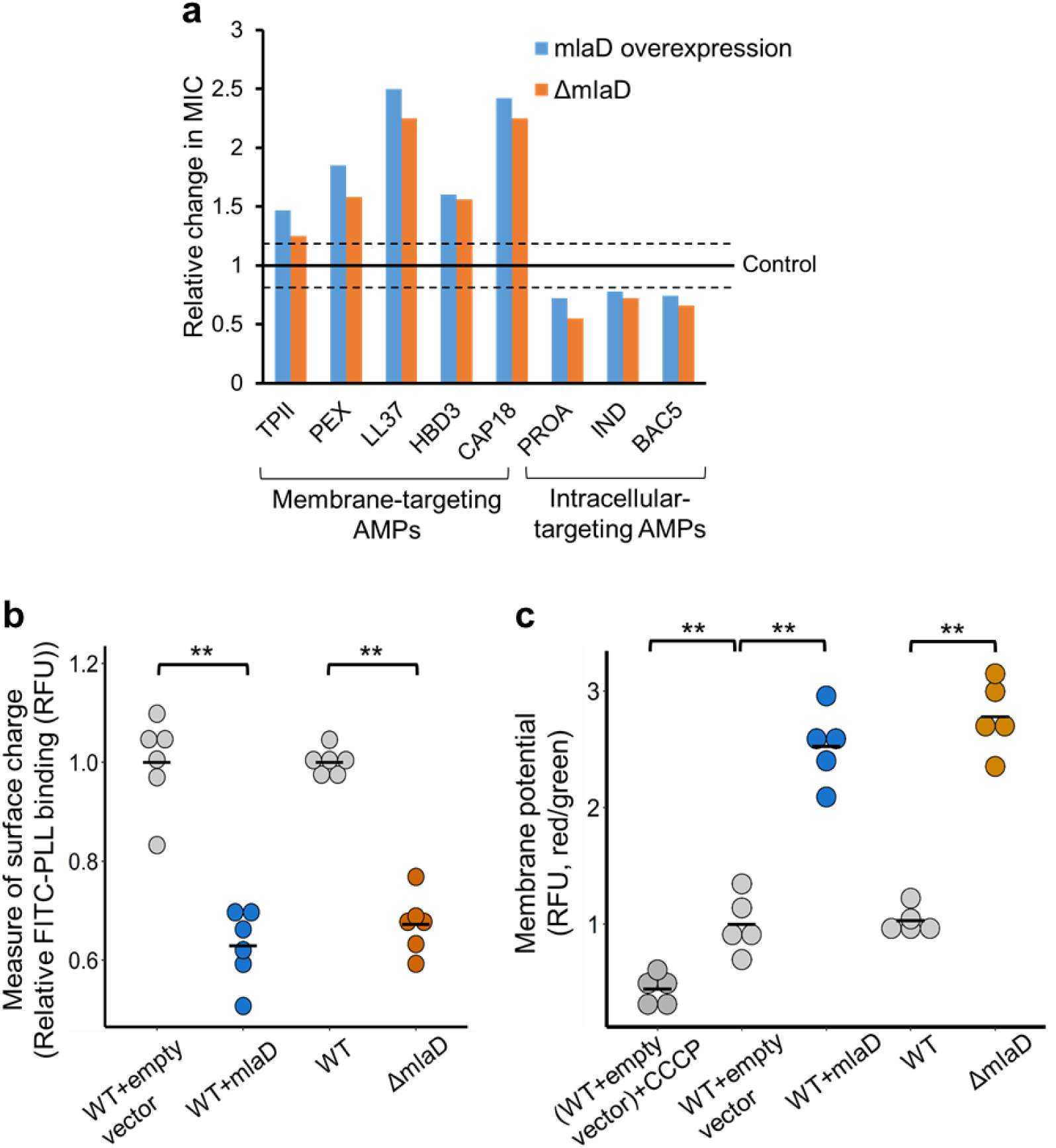
Mutation in *mlaD* influences AMP susceptibilities through antagonistic mutational effects. **a,** Relative change in MICs of the mlaD overexpression and deletion strains (*ΔmlaD*) to a representative set of membrane-targeting and intracellular-targeting AMPs. MICs were compared to corresponding wild-type control strains (see Supplementary Figures 8 and 9). Dashed lines represent previously defined cut-offs for resistance (≥1.2 x MIC of the control) and sensitivity (≤0.8 x MIC of the control)^27^. **b,** Decreased net negative surface charge of the *mlaD* overexpression and deletion strains. Significant differences: ** *P* = 0.0021 and *P* = 0.0021 for WT+empty vector vs overexpression and WT vs deletion strain, respectively, from two-sided Mann–Whitney U test, *n* = 6 biological replicates for each genotype. Charge measurement was done using FITC-labelled poly-L-lysine (FITC-PLL) assay where the fluorescence signal is proportional to the binding of the FITC-PLL molecules. A lower binding of FITC-PLL indicates a less net negative surface charge of the outer bacterial membrane (see Methods). **c,** Increased membrane potentials of the *mlaD* overexpression and deletion strains. Significant differences: ** *P* = 0.007, *P* = 0.0079 and *P* = 0.0079 for WT+empty vector CCCP control vs WT+empty vector, WT+empty vector vs WT+mlaD overexpression and WT vs. *ΔmlaD,* respectively, two-sided Mann–Whitney U test, *n* = 5 biological replicates for each genotype. Relative membrane potential was measured by determining relative fluorescence (RFU) using a carbocyanine dye DiOC2(3) assay (see Methods). Red/green ratios were calculated using population mean fluorescence intensities. WT *E. coli* carrying the empty vector treated with CCCP was used as an experimental control for diminished membrane potential. Raw data is in Supplementary Figure 13.

It has been observed that *mlaD* deletion alters the membrane composition by leading to the accumulation of phospholipids in the outer leaflet of the bacterial outer membrane^39^. A change in membrane composition can alter the net negative surface charge of the cell^3^, which in turn strongly influences AMP susceptibility^1^. Thus, we hypothesized that depletion of functional MlaD decreases susceptibility to membrane-targeting AMPs by decreasing the net negative surface charge of the cell. On the other hand, membrane properties can also have an effect on membrane potential^45^. As the uptake of certain intracellular-targeting AMPs, for example, PROA and IND, are driven by membrane potential^46,47^, we posited that such an effect could underlie the observed collateral sensitivity interactions. To test this, we measured the net negative surface charge and the membrane potential of the MlaD overexpression and deletion strains (see Methods). Reassuringly, both overexpressing and deleting *mlaD* resulted in a significantly decreased negative surface charge (Figure 5b) and in increased membrane potential (Figure 5c).

### Chemical-genetic profiles predict the cross-resistance spectrum of human-B-defensin-3

It has recently been proposed that chemical-genetics could be employed to infer whether resistance evolution to an antimicrobial agent would lead to cross-resistance to another agent^12^. In particular, a high overlap in the latent resistomes may indicate the emergence of cross-resistance during evolution in nature or in the laboratory. If so, the extent of cross-resistance between AMPs may show a pattern that follows the observed clusters in the chemical-genetic map.

To test this notion, we performed an adaptive laboratory evolution experiment against human-B-defensin-3 (HBD-3). We have chosen this AMP due to its relevance as a human host-defense peptide, and because its chemical-genetic profile is similar to AMPs belonging to cluster C2, but markedly different from the rest of AMPs (Figure 1c and 2a). Ten parallel *E. coli* cell populations were propagated under increasing concentration of HBD-3 for approximately 120 bacterial generations (see Methods). An approximately 10-fold increase in minimum inhibitory concentration (MIC) was observed in the evolved lineages (Supplementary Figure 10). Out of the 10 evolved lines, 4 were subjected to whole-genome sequencing to identify the mutations underlying elevated AMP resistance. Genome sequence analysis revealed a total of 27 unique mutational events (including large deletions, Supplementary Table 9).

We then measured how the susceptibility of the four evolved lines changed to a set of 10 AMPs representing the four major chemical-genetic clusters (C1 to C4). We found that high levels of cross-resistance occurred only to AMPs that were clustered together with HBD-3 in the chemical-genetic map (C2), while cross-resistance to AMPs belonging to the other three clusters (C1, C3, C4) were rare (Figure 6a). These results demonstrate that clustering of the chemical-genetic profiles predicted the observed cross-resistance patterns of the HBD-3-adapted lines.

**Figure 6.**
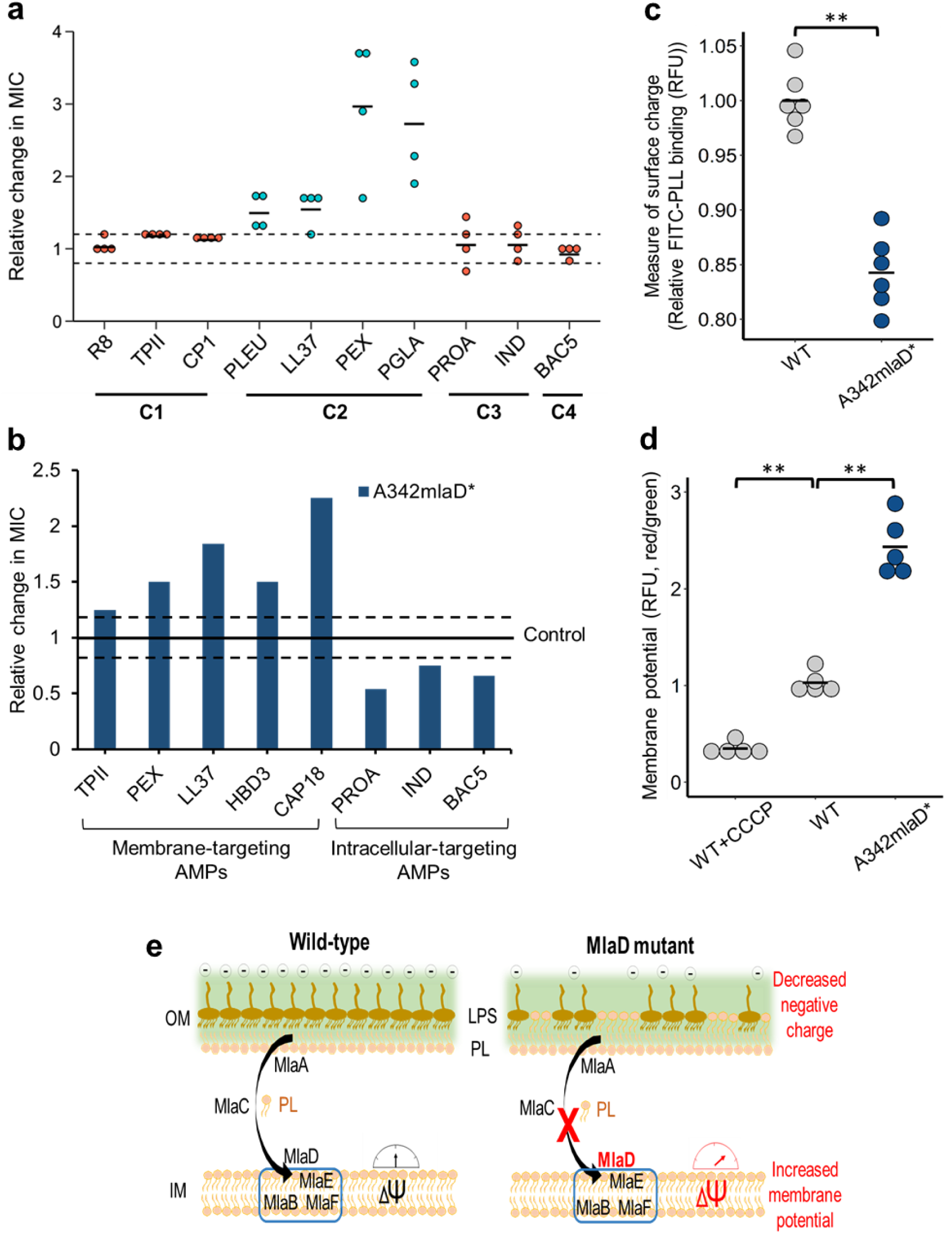
Chemical-genetic profiles predict the cross-resistance spectrum of HBD-3. a, HBD-3-evolved lines showed cross-resistance almost exclusively to AMPs from the same chemical-genetic cluster (*P* = 0.0003, two-sided Fisher’s exact test, *n* = 16 and 24 for AMPs from C2 (blue points) and from the rest of the clusters (red points), respectively). Relative MICs were determined for each of the 4 sequenced evolved lines by comparing their MICs to that of the ancestral cell line (control). Dashed lines are previously defined cut-off values for cross-resistance (≥1.2 x MIC of the control) and collateral sensitivity (≤0.8 x MIC of the control)^27^. Data is provided in Supplementary Table 11. **b,** Relative change in MICs of the A342mlaD* mutant strain to a representative set of AMPs. MICs were compared to corresponding wild-type control strain (see Supplementary Figures 12). **c,** Decreased net negative surface charge of the A342mlaD* mutant strain (significant difference: ** *P* = 0.0021, two-sided Mann–Whitney U test, *n* = 6). **d,** Increased membrane potential of the A342mlaD* mutant strain (significance differences: ** *P* = 0.0079 and *P* = 0.0079 for WT+CCCP control vs WT strain and WT vs A342mlaD* mutant, two-sided Mann– Whitney U test, *n* = 5). WT *E. coli* treated with CCCP was used as a control for diminished membrane potential. **e,** Proposed molecular mechanism for the collateral sensitivity caused by the perturbation of *mlaD.* Accumulation of phospholipids in the outer membrane upon depletion of functional *mlaD* results in a decreased net negative surface charge which causes weaker electrostatic interaction between the bacterial membrane and the AMPs thereby providing resistance to membrane-targeting AMPs. At the same time, this mutation also increases membrane potential which drives the cellular uptake of certain intracellular-targeting peptides. Abbreviations: OM outer membrane, IM inner membrane, PL phospholipid, LPS lipopolysaccharides, ΔΨ membrane potential.

Next, we interrogated if the chemical-genetic profiles provide an insight into the molecular mechanisms underlying the cross-resistance patterns of the HBD-3-evolved lines. To this end, we first analyzed the function of the mutated genes. Phospholipid and LPS-related genes were overrepresented among the mutated genes (Supplementary Figure 11a). Specifically, three HBD3-evolved lines carried mutations in genes involved in retrograde phospholipid transport (Mla pathway, see Supplementary Table 9). As overexpression of genes in this pathway induced resistant chemical-genetic interactions to membrane-targeting peptides only (Supplementary Figure 7 and 11b), we hypothesized that the mutations in the phospholipid transport genes contribute to the observed cross-resistance patterns in the evolved lines (Figure 6a).

To test this hypothesis, we reconstructed an adaptive mutation (A342mlaD*) which was identified in a HBD-3-evolved line by inserting it into the *mlaD* gene of wild-type *E. coli*. Then, we measured the susceptibility of this mutant to a selected set of membrane-targeting (C1-C2) and intracellular-targeting AMPs (C3-C4). As expected, the strain carrying the A342mlaD* mutation showed a decreased susceptibility to membrane-targeting AMPs and an increased susceptibility to intracellular-targeting ones, similarly to the MlaD overexpression strain (Figure 6b, for MIC curves, see Supplementary Figure 12). In agreement with these findings, measuring the net negative surface charge and the membrane potential of the mutant strain confirmed the same mutational effects as in the case of the *mlaD* overexpression and deletion strains (Figure 6c and 6d).

In sum, the chemical-genetic profiles predicted the observed cross-resistance patterns of the HBD-3-evolved lines and illuminated the mechanistic basis thereof.

## Discussion

This work systematically mapped the genetic determinants of AMP resistance by chemical-genetic profiling in *E. coli* (Figure 1). We report that AMP resistance is influenced by a large set of functionally diverse genes, and yet these genes overlap only to a limited extent between AMPs. Specifically, clustering of the chemical-genetic profiles revealed that the modes of action of the AMPs largely define the gene sets that influence bacterial susceptibility against them (Figure 2 and 3). Additionally, antagonistic mutational effects are frequent between AMPs that disrupt the bacterial membrane versus those that act on intracellular targets (Figures 4 and 5). Finally, by applying adaptive laboratory evolution in the presence of HBD-3, a human AMP targeting the bacterial membrane, we show that the clustering of the chemical-genetic profiles predicts the cross-resistance spectra of the evolved *E. coli* lines across different groups of AMPs (Figure 6). This indicates that cross-resistance between AMPs are shaped by the overlap in chemical-genetic profiles.

The results presented in this study have important implications for the development of AMP-based therapies. Previous works reported several instances of cross-resistance interactions between membrane-targeting peptides, however, the potential for cross-resistance across AMPs with different modes of action has remained poorly understood. Specifically, while cross-resistance between host and therapeutic AMPs is certainly a realistic danger, not all AMPs are equally prone to cross-resistance. Given the immense diversity of AMPs with major differences in physicochemical properties and resistance mechanisms, we propose that carefully chosen therapeutic candidates could mitigate the risk of cross-resistance with specific human host-defense peptides. From our screen, proline-rich AMPs are the best candidates in this respect, supporting the considerable effort that has already been taken into the clinical development of proline-rich AMP-based therapeutic applications^48,49^. Additionally, a distinct group of membrane-targeting AMPs (R8, TPII and CP1) appear to be less prone to cross-resistance to the investigated human host-defense AMPs. Clearly, this work made the first step in this direction and future studies should explore these possibilities. Specifically, cross-resistance patterns of proline-rich AMPs in human saliva and synthetic AMPs should also be considered^50^.

The large diversity of genes that influence AMP resistance upon overexpression indicates that bacterial susceptibility to AMPs is coupled to the general physiology of the bacterial cell, and in particular to alterations in membrane composition. This idea also provides an explanation to a recent finding that antibiotic resistance mutations in membrane proteins frequently induce collateral sensitivity to AMPs through pleiotropic side effects that alter membrane composition^27^. Indeed, the overrepresentation of collateral sensitivity interactions among AMP resistance determinants implies that evolving AMP resistance requires the optimization of many traits simultaneously. As a consequence, bacterial cells potentially harbor a large mutational target to alter AMP resistance, however, such mutations often have negative trade-offs with other cellular traits.

Collateral sensitivity between AMPs is best exemplified by the Mla pathway. Several studies have reported the importance of Mla pathway in bacterial pathogenesis and virulence^40–42^. For example, loss-of-function mutations in Mla pathway in *Haemophilus influenzae* increased the accumulation of phospholipids in the outer membrane, which mediated sensitivity to human serum^40^. Here, we demonstrated that a loss-of-function mutation in *mlaD* decreases the net negative surface charge of the bacterial membrane and, eventually, causes a somewhat increased resistance to human membrane-targeting AMPs (Figure 5a,b, 6b,c), and an elevated susceptibility to intracellular-targeting AMPs (Figure 6e). Together, our work indicates that a trade-off between membrane surface charge and membrane potential underlie collateral sensitivity interactions between membrane-targeting and intracellular-targeting AMPs upon perturbing the Mla pathway. We speculate that this trade-off could contribute to the observed variation in the expression level of Mla pathway proteins among clinical isolates of *H. influenzae*^40^.

Our results also have implications to an important but unresolved issue: why have natural AMPs that are part of the human innate immune system remained effective for millions of years without detectable resistance in several bacterial species? One possibility, supported by our work, is that bacteria may have difficulty to evolve resistance to the combination of multiple defense peptides deployed by the immune system due to negative trade-offs between them. We do not wish to claim, however, that AMPs in clinical use would generally be resistance-free. Rather, these properties of the AMPs could be beneficial for the development of combination therapies involving AMPs in combination with antibiotics and human host peptides.

## Materials and methods

### Media, bacterial strains and antimicrobial peptides

Experiments with AMPs were conducted in minimal salts (MS) medium supplemented with MgSO_4_ (0.1 mM), FeCl_3_ (0.54 μg/ml), thiamin (1 μg/ml), casamino acids (0.2%) and glucose (0.2%). Luria-Bertani (LB) medium contained tryptone (0.1%), yeast extract (0.05%), and NaCl (0.05%). All components were purchased from Sigma-Aldrich. To increase the dosage of each *Escherichia coli* gene for the chemical-genetic screen, we used the *E. coli* K-12 Open Reading Frame Archive library (ASKA)^23^ in *Escherichia coli* K12 BW25113 cells. AMPs were custom synthesized by ProteoGenix, except for protamine and polymyxin B, which were purchased from Sigma-Aldrich. AMP solutions were prepared in sterile water and stored at −80°C until further use.

### Plasmid DNA preparation and purification

Bacterial cells harbouring the ASKA plasmids were grown overnight in LB medium supplemented with chloramphenicol (20 µg/ml). Cells were harvested by centrifugation. Plasmid DNA isolation was performed using innuPREP plasmid mini Kit (Analytik Jena AG) according to the manufacturer’s instructions. To remove the genomic DNA contamination, the isolated plasmid DNA samples were digested overnight with Lambda exonuclease and exonuclease I (Fermentas) at 37°C. The digested plasmid DNA samples were purified with DNA Clean & Concentrator™(Zymo) kit according to the manufacturer’s instructions.

### Chemical-genetic profiling

We carried out chemical-genetic profiling to determine the impact of the overexpression of each *E. coli* ORF on bacterial susceptibility to each of the 15 different AMPs. To this end, we used the ASKA plasmid library (GFP minus) where each *E. coli* ORF is cloned into a high copy number expression plasmid (pCA24N-ORFGFP(-)). Prior to screening, this plasmid collection was pooled and transformed into *E. coli* K12 BW25113 strain as described before^51^. To obtain a negative control strain not having any overexpressed gene, the plasmid without a cloned ORF (pCA24N-noORF) was also transformed into the same *E. coli* strain. Then, on the pooled collection, we applied a previously reported competitive growth assay^27^. Specifically, the pooled overexpression library and the control strain were grown in parallel in MS medium supplemented with 20 µg/ml chloramphenicol and the overexpression was induced by 100 µM isopropyl-ß-D-thiogalactopyranoside (IPTG). After 1h induction, ∼5 x 10^5^ bacterial cells from the library were inoculated into each well of a 96-well microtiter plate containing a concentration gradient of an AMP in the MS medium supplemented with 20 µg/ml chloramphenicol and 100 µM IPTG. At the same time, both the library and the control strain with the empty plasmid were grown in the absence of any AMPs. We took special care to grow both of these samples in the exact same conditions as the samples in the presence of AMPs. Bacterial growth was monitored in a microplate reader (Biotek Synergy 2) for 24 h. At the end of the exponential growth phase, we selected those wells in which the doubling time of the cell population was increased by 2-fold. Then, from these wells, cells were split into four equal proportion and each was transferred into 20 mL of MS medium supplemented with the corresponding AMP in four different concentrations in the range that slowed down growth by two-fold in the microtitre plate. Then, following exponential growth, out of the four 20 mL cultures those that showed again a two-fold increase in doubling time were selected for further analysis. The rationale for this 2-step process was to maintain competition in exponential phase for 12 generations of growth, efficiently control the growth rate in a reproducible manner and obtain the plasmid pool with standard DNA isolation protocol (innuPREP plasmid mini Kit, Analytik Jena AG) in a yield that is enough for the downstream analysis. Each of the selected plasmid samples was digested overnight with a mixture of lambda exonuclease and exonuclease I (Fermentas) at 37°C to remove the genomic DNA background. The digested plasmid DNA samples were purified with DNA Clean & Concentrator™(Zymo) kit according to the manufacturer’s instructions. This protocol was carried out in two biological replicates for each AMP treatment. In the case of the untreated sample (in the absence of AMP), we had five replicates. *E. coli* BW25113 strain carrying the empty vector was used as a negative control to measure read counts originating from genomic DNA contamination during plasmid preparation (background).

### Deep sequencing of plasmid pool

The cleaned plasmid samples were sequenced with the SOLiD next-generation sequencing system (Life Technologies) and the relative abundance of each plasmid was determined as described previously^27,51^. Briefly, the isolated plasmid pool samples were fragmented and subjected to library preparation. Library preparation and sequencing was performed using the dedicated kits and the SOLiD4 sequencer (Life Technologies), respectively. For each sample, 20-25 million of 50 nucleotide long reads were generated. Primary data analysis was carried out with software provided by the supplier (base-calling). The 50 nucleotide long reads were analyzed, quality values for each nucleotide were determined using the CLC Bio Genomics Workbench 4.6 program.

### Data analysis of chemical-genetic screen

Raw sequence data processing and mapping onto *E. coli* ORFs were carried out as described previously^27^. Raw sequence data were also mapped to the plasmid backbone. In order to make the mapped read counts comparable between the different samples, we carried out the following data processing workflow based on established protocols^52,53^, using a custom-made R script. The extra read counts deriving from genomic DNA contamination (background) were estimated by assuming that the reads mapping to the unit length of the plasmid and the ORFs should have a ratio of 1:1. The total extra read count estimated thereof was partitioned among the ORFs based on their background frequency (that is, their relative frequency obtained from the experiment involving the empty plasmid). Next, these ORF-specific backgrounds were subtracted from the read counts. Then, a loglinear transformation was carried out on the background-corrected relative read counts. Compared to the canonical logarithmic transformation, this transformation has the advantage of avoiding the inflation of data variance for ORFs with very low read counts^54^. The transformed relative read counts showed bimodal distributions (Supplementary Figure 14). The lower mode of the distribution corresponds to ORFs that were not present in the sample. The upper mode represents those ORFs whose growth was unaffected by overexpression (i.e. no fitness effect). To make different samples comparable, the two modes of the distribution of each sample were set to two predefined values. These values were chosen such that the original scale of the data was retained. In order to align the modes between samples, we introduced two normalization steps: one before and one after loglinear transformation. The first normalization step identified the lower mode corresponding to the absent strains and added a constant to shift the lower mode to zero. Next, we performed the loglinear transformation step described above. The second normalization step was a linear transformation moving the upper mode to a higher predefined value. Following these normalization steps, genes that were close to the lower mode in the untreated samples were discarded from the analysis as these represent strains that displayed poor growth even in the absence of drug treatment (that is, AMP sensitivity could not be reliably detected). A differential growth score (i.e. fold-change) was calculated for each gene as the ratio of the normalized relative read counts in treated and non-treated samples at the end of the competition. Fold-change values of biological replicate experiments were averaged. Genes that showed at least 2-fold lower and higher relative abundance at the end of the competition upon AMP treatment were considered as sensitive and resistant genes, respectively.

### Cluster analysis of chemical-genetic profiles

To group AMPs with similar chemical-genetic profiles, we employed an ensemble clustering algorithm that combines multiple clustering results to obtain a robust clustering^55^. A combination of diverse clustering results based on perturbing the input data and clustering parameters is known to yield a more robust grouping of data points than that obtained from a single clustering result.

As a first step, we removed genes that did not show AMP-specific phenotypes across treatments since these genes would be uninformative for clustering. To this end, we retained only those genes that showed significant differences in their fold-change values between AMPs compared to their variances across replicate measurements within AMPs as assessed by F-tests (p<0.01). This resulted in a set of 2146 genes kept for clustering. Next, we employed a distance metric, normalized variation of information, to measure distances between AMP chemical-genetic profiles. The normalized variation of information is closely related to mutual information but has the advantage of being a true distance metric. Importantly, normalized variation of information gives more weight to rare overlaps of resistance/sensitivity phenotypes between AMPs, unlike the commonly used Euclidean distance. Normalized variation of information (NVI) between AMP pairs was calculated as follows: NVI = (H I) / H where H is the entropy and I is the mutual information.

Based on this distance measure, we then generated 75,000 clusters of AMPs by perturbing both the AMP profile data and the clustering parameters. The AMP profile data was perturbed by resampling the gene set with replacement and by randomly selecting a single chemical-genetic profile among the multiple biological replicates available for each AMP. We used hierarchical clustering and varied both the algorithms (Ward, single-linkage, complete-linkage and average-linkage) and the number of clusters defined (k = 2…6). Results of the 75,000 clusters were summarized in a consensus, which contains, for each pair of AMP, the number of times that two AMPs cluster together across all of the clustering results. Finally, we clustered this consensus matrix using hierarchical clustering and complete linkage and plotted the result as a heatmap.

### Construction of hypomorphic alleles for chemical-genetic screening

A total of 279 essential gene hypomorphs (with reduced protein expression) were constructed essentially as previously described^20,21^. Briefly, as with the mRNA perturbation by DAmP (decreased abundance by mRNA perturbation) alleles in yeast^56^, we created an essential gene hypomorphic mutation by introducing a kanamycin (Kan^R^) marked C-terminal sequential peptide affinity fusion tag, engineered by homologous recombination into each essential gene^57^. The tag perturbs the 3’ end of the expressed mRNA of the essential proteins, when combined with environmental/chemical stressors, or other mutations by destabilizing the transcript abundance. A subset of these hypomorphic alleles that we used^21,22,58,59^ or shared with others^16^ have revealed functionally informative genegene, and gene-environment or drug-gene interactions.

Analogous to our *E. coli* synthetic genetic array approach^58^, our chemical-genetics screening strategy involves robotic pinning of each Kan^R^ marked single essential gene hypomorph arrayed in 384 colony format on Luria Broth (LB) medium, in quadruplicate, onto the minimal medium containing AMPs under a selected concentration, in two replicates, generating eight replicates in total for each essential gene hypomorph. The sub-inhibitory concentration was chosen based on 50% growth inhibition of wild-type cells using a serial dilution. In parallel, we also prepared two replicates of control plates containing arrayed essential gene hypomorphic strains pinned onto minimal media without AMPs. After incubation at 32°C for 20 h, the plates (with and without AMPs) were digitally imaged and colony sizes were extracted from the imaged plates using an adapted version of the gitter toolbox^60^. The resulting raw colony size (proxy for cell growth) from each screen, with and without AMP, was normalized using SGAtools suite^61^, with default parameters. The normalized colony sizes from the AMP plate was subtracted from their corresponding colony screened without AMP to estimate the final hypomorphic-strain fitness score (sensitive or resistant), which is as an average of all eight replicate measurements recorded for each hypomorphic allele.

### Physicochemical properties of AMPs

Protein amino acid frequencies were counted with an inhouse perl script. Isoelectric point, hydrophobicity, hydrophobic moment, net charge and membrane surface was calculated with the peptides R package, version 2.4^62^. The ExPasy Prot Param tool was used for calculating molecular weight and peptide length^63^.

### Differentiation between AMP clusters based on physicochemical parameters

Logistic regression framework was used with two parameters to infer differences between C1 and C2 clusters in the peptides physicochemical properties. Area under the receiver operating characteristic curve (ROC) was used to establish model accuracies and rank parameter pairs using the caret (v6.0-80) and e1071 (v1.7-0) R packages. For a global analysis of cluster properties, principal component analysis was applied to all the peptide physicochemical properties with centering and scaling the data using the princomp R package. All calculations were done in R version 3.5.0 in Rstudio version 1.1.447^64,65^.

### Calculating the overlap in resistance / sensitivity gene sets between AMPs

To calculate the extent to which the resistance- and sensitivity-conferring genes are shared between pairs of AMPs, we used a modified version of the Jaccard index that takes into account measurement noise. Specifically, for each pair of AMP, we calculated the Jaccard index of overlap between their sets of resistance genes and performed a correction by dividing this value by the average Jaccard index of overlap between replicate screens of the same AMPs. Thus, a corrected Jaccard index value of 1 between two AMPs indicates that the set of resistance genes overlap as much as that of two replicate screens.

### Enrichment of collateral sensitivity interactions between AMP pairs

We calculated the overrepresentation of collateral sensitivity-inducing genes for each AMP pair over random expectation using data from our overexpression screen. Random expectation was calculated using the number of resistance and sensitive genes for each AMP. Enrichment ratio (r) of collateral sensitivity-inducing genes for each AMP pair was calculated as follows:

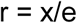

where:
x- actual frequency of the genes showing collateral sensitivity interactions between AMP pair
e- expected frequency (based on marginal probability) of the genes showing collateral sensitivity interactions between AMP pair. Expected frequency (e) was calculated as follows:

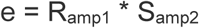

where:

R_amp1_ = relative frequency of genes showing resistance to AMP1 out of all ∼4400 genes screened

S_amp2_ = relative frequency of genes showing sensitivity to AMP2 out of all the ∼4400 genes screened.

### Gene-ontology (GO) enrichment analysis

To determine which Gene-ontology (GO) terms are significantly enriched in the resistant and sensitive genes, we employed the Biological Networks Gene Ontology tool (BiNGO)^66^. The selection of GO reference set was based on the EcoGene database^67^. The Benjamini-Hochberg FDR (FDR cutoff = 0.05) was used for multiple-testing correction^68^. GO categories showing FDR-corrected P-values <0.05 were considered statistically significant. Detailed information about the significantly enriched GO categories is provided in Supplementary Table 7.

We calculated the enrichment of phospholipid and lipopolysaccharide transport/binding functions among the set of *E. coli* genes that were mutated in HBD-3-adapted lines. The same enrichment analysis was also carried out for those genes that showed collateral sensitivity. Genes related to phospholipid and lipopolysaccharide transport/binding function were selected from a previous study^27^.

### Determination of minimum inhibitory concentration (MIC)

Minimum inhibitory concentrations (MIC) were determined with a standard serial broth dilution technique^69^. Briefly, from a stock solution of an AMP, 12-steps serial dilution was prepared in fresh MS medium in 96-well microtiter plates. Each AMP was represented in 11 different concentrations (3 wells/AMP concentration/strain). Three wells contained only medium to check the growth in the absence of AMP. After overnight growth in MS medium supplemented with chloramphenicol, bacterial strains were diluted 20-fold into fresh MS medium and grown until the cell density reached OD_600_ ∼1. Cells were induced by 100 µM of IPTG and incubated for 1 h at 30°C with continuous shaking at 300 rpm. Following incubation, approximately half-million cells were inoculated into the wells of the 96-well microtiter plate with a 96-pin replicator. We used three independent replicates for each strain and the corresponding control. Two rows in the 96-well plate contained only MS medium in order to obtain the background OD value of the medium. Plates were incubated at 30°C with continuous shaking at 300?rpm. After 20-24 h of incubation, OD_600_ values were measured in a microplate reader (Biotek Synergy 2). After background subtraction, MIC was determined as the lowest concentration of AMP where the OD_600_ values were less than 0.05.

### Membrane surface charge measurement

To evaluate bacterial surface charge, we performed a fluorescein isothiocyanate-labeled poly-L-lysine (FITC-PLL) (Sigma) binding assay. In brief, FITC-PLL is a polycationic molecule that binds to anionic lipid membrane in a charge-dependent manner and is used to investigate the interaction between cationic peptides and charged lipid bilayer membranes^70,71^. The assay was performed as previously described^27,70^. Briefly, bacterial cells were grown overnight in MS medium, centrifuged and washed twice with 1X PBS buffer (pH 7.4). The washed bacterial cells were re-suspended in 1XPBS buffer to a final OD_600_ of 0.1. A freshly prepared FITC-PLL solution was added to the bacterial suspension at a final concentration of 6.5 µg/ml. The suspension was incubated at room temperature for 10 minutes, and pelleted by centrifugation. The remaining amount of FITC-PLL in the supernatant was determined fluorometrically (excitation at 500 nm and emission at 530 nm) with or without bacterial exposure. The quantity of bound molecules was calculated from the difference between these values. A lower binding of FITC-PLL indicates a less net negative surface charge of the outer bacterial membrane.

### Membrane potential measurement

A previously described protocol^37^ was used to determine the change in transmembrane potential for *mlaD* overexpression, *mlaD* knockout and A342*mlaD*\* mutant strains in comparison to their control strain. Transmembrane potential (ΔΨ) was measured using the BacLight™ Bacterial Membrane Potential Kit (Invitrogen). In this assay, a fluorescent membrane potential indicator dye emits green fluorescence in all bacterial cells and the emission shifts to red in the cells that maintain a high membrane potential. In this way, the ratio of red/green fluorescence provides a measure of membrane potential. Prior to the measurement bacterial cells were grown overnight in MS medium at 30°C. The overnight cultures were diluted into fresh MS medium and grown until cell density reached OD_600_ 0.5-0.6. The grown cultures were diluted to 10^6^ cells/mL in filtered PBS buffer. Then, 5 µl of 3 mM DiOC_2_(3) was added to each sample tube containing 500 µl of bacterial suspension and incubated for 20 minutes at room temperature. Following incubation, red to green fluorescence values of the samples were measured using Fluorescence Activated Cell Sorter (BD Facscalibur) according to the instructions of the kit’s manufacturer. Fluorescence values were calculated relative to the control strain. Control populations treated with cyanide-m-chlorophenylhydrazone (CCCP, a chemical inhibitor of proton motive force) were used as an experimental control.

### Experimental evolution of resistance

Adaptive laboratory evolution experiment was performed using a previously established automated evolution experiment protocol^27,72^. Briefly, starting from a single clone of *E. coli* BW25113, 10 parallel cultures were grown in the presence of sub-inhibitory concentration of HBD-3. A chess-board layout was used on the plate to monitor cross-contamination events. Each culture was allowed to grow for 24 h. Following incubation 20 µl of the grown culture was transferred to four independent wells containing fresh MS medium and increasing dosages of HBD-3 (0.5x, 1x, 1.5x and 2.5x the concentration of the previous step). Prior to each transfer, cell growth was monitored by measuring the optical density at 600 nm. Only populations of the highest drug concentration that reached OD_600_ > 0.2 were selected for further evolution. Accordingly, only the population of one of the four wells was retained for each independently evolving lineage. This protocol was designed to avoid population extinction and to ensure that populations with the highest level of resistance were propagated further during evolution. The experimental evolution was maintained through 20 transfers. Following that, MIC values of HBD-3 were determined for all evolved lineages.

### Whole-genome sequencing of HBD-3-evolved lines

To identify potential mechanisms conferring resistance to the human beta-defensin-3 (HBD-3), 4 out of 10 adapted lines were subjected to whole-genome sequencing. Isolation of bacterial genomic DNA was performed using Sigma GenElute™ Bacterial Genomic DNA Kit and quantified using Qubit dsDNA BR assay in a Qubit 2.0 fluorometer (Invitrogen). 200 ng of genomic DNA was fragmented in a Covaris M220 focused-ultrasonicator (peak power: 55W, duty factor: 20%, 200 cycles/burst, duration: 45 sec) using Covaris AFA screw cap fiber microTUBEs. Fragment size distribution was analyzed by capillary gel electrophoresis using Agilent High Sensitivity DNA kit in a Bioanalyzer 2100 instrument (Agilent) then indexed sequencing libraries were prepared using TruSeq Nano DNA LT kit (Illumina) following the manufacturer’s protocol. This, in short, includes end repair of DNA fragments, fragment size selection, ligation of indexed adapters and library enrichment with limited-cycle PCR. Sequencing libraries were validated (library sizes determined) using Agilent High Sensitivity DNA kit in a Bioanalyzer 2100 instrument then quantitated using qPCR based NEBNext Library Quant Kit for Illumina (New England Biolabs) with a Piko-Real Real-Time PCR System (Thermo Fisher Scientific) and diluted to 4 nM concentration. Groups of 12 indexed libraries were pooled, denatured with 0.1 N NaOH and after dilution loaded in a MiSeq Reagent kit V2-500 (Illumina) at 8 pM concentration. 2X250 bp pair-end sequencing was done with an Illumina MiSeq sequencer, primary sequence analysis was done on BaseSpace cloud computing environment with GenerateFASTQ 2.20.2 workflow. Paired end sequencing data were exported in FASTQ file format. The reads were trimmed using Trim Galore and cutadapt to remove adapters and bases where the PHRED quality value was less than 20. Trimmed sequences were removed if they became shorter than 150 bases. FASTQC program (https://www.bioinformatics.babraham.ac.uk/projects/fastqc/) was used to evaluate the qualities of original and trimmed reads. The Breseq program was used with default parameters for all samples^73^. The gdtools package was used for annotating the effects of mutations and comparing multiple samples. The genbank formatted reference genome BW25113.gb was used as a reference genome in the analysis.

### MlaD knockout and A342mlaD* mutant construction

A *mlaD* knockout strain of *E. coli* BW25113 carrying a kanamycin resistance cassette in the position of the gene was selected from the KEIO collection^74^. The resistance marker was removed using plasmid borne (pFT-A) expression of FLP recombinase leading to excision of the kanamycin resistance cassette^75^. Cassette excision was verified by a polymerase chain reaction using the primers mlaD_del_ver_Fw (5’TCACGGTGACGTGGATTTC) and mlaD_del_ver_Rev (5’GCCTCGTCCATCAGCTTATAC). The identified single nucleotide deletion at position 342 within the *mlaD* gene was constructed in *E. coli* BW25113. A well-established recombineering-based method employing pORTMAGE-2^76^ was used to introduce the specific deletion. The mlaD_a342del ssDNA oligo (5’-A*T*TG TATCGCCATCCTTCAGGATAGCAGTCCCCAGTTCCGGGTCTTCAAAACCGACGTTAATGCCAGATATTGTTCCCCCAGCAGGCC*G*G) was employed to introduce the specific deletion (where * denotes a phosphorothioate bond). Cells were then screened using allele-specific PCR using the mlaD_ASP (5’GGAACAATATCTGGCATTAACG) and mlaD_Rev (5’GTCATCGCCTTTACTACC) primer pair. Candidates were finally verified by Sanger sequencing using the mlaD_Seq (5’-TACTGAACCGACCTACAC) primer paired with mlaD_Rev.

## Acknowledgements

We thank Roland Tengölics for his helpful discussion. This work was supported by the ‘Lendület’ programme of the Hungarian Academy of Sciences (B.P. and C.P.), the Wellcome Trust (B.P.), The European Research Council H2020-ERC-2014-CoG 648364Resistance Evolution (C.P.), GINOP2.3.2-15-2016-00014 (EVOMER, C.P. and B.P.), GINOP-2.3.2-15-2016-00020 (MolMedEx TUMORDNS, C.P.) and GINOP-2.3.2-15-2016-00026 (iChamber, B.P.), National Research, Development and Innovation Office, Hungary NKFIH grant K120220 (B.K.), NKFIH grant FK124254 (O.M.) and NKFIH grant KH125616 (B.P.), Natural Sciences and Engineering Research Discovery Grant 20234-2012 and Canadian Institutes of Health Research (CIHR) project grant 148831 (M.B). A.Ga. is supported by a CIHR Postdoctoral Fellowship. L.B. was supported by grant NKFI-112294. B.K. holds a Bolyai Janos Scholarship and supported by the UNKP-18-4 New National Excellence Program of the Ministry of Human Capacities.

## Author contributions

B.K., C.P. and B.P. conceived the project, B.K., P.K.J., G.F., C.P. and B.P. planned experiments and data analyses. P.K.J. and M.S. performed most experiments. R.S., L.D., A.M. and V.L. carried out laboratory evolution. I.N. was responsible to Solid and Illumina sequencing. A.F. and L.B. performed whole-genome sequencing of HBD-3-evolved lines. B.C. carried out mutagenesis. B.K., P.K.J., F.G., O.M., E.A. analyzed the experimental data. P.K.J., G.F. and A.G. carried out bioinformatic analyses. A.H., A.Ga. and S.K. created all essential hypomorphic alleles, and performed the chemical-genetic screening. S.P quantified the colony growth fitness of the hypomorphs, and analyzed the data with input from M.B. B.K., P.K.J., C.P., and B.P. wrote the manuscript.

## Data and code availability

All data generated or analysed during this study are present in this article and its Supplementary Information files. For each figure, the availability of the analysed data is indicated in the figure legend. The Illumina and SOLiD sequencing data for the chemical-genetic screen will be available in the NCBI Sequence Read Archive, with SRA accession number XX. Any additional data can be requested from the corresponding author.

All scripts and other files needed to reproduce our analyses will be available at https://github.com before publication.

