## Supplementary information for "Chemical-genetic profiling reveals cross-resistance and collateral sensitivity between antimicrobial peptides"

[5]Present address: Department of Microbiology and Immunology, University of California, San Francisco, CA 94143, USA.

[7]Present address: Faculty of Biology, Technion – Israel Institute of Technology, Haifa, Israel

### Supplementary Figures

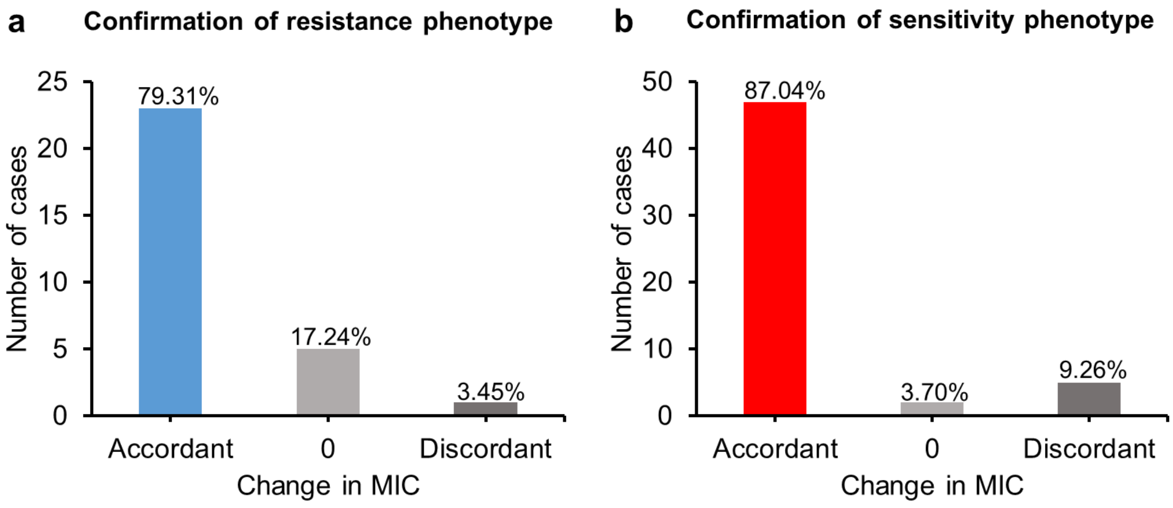


**Supplementary Figure 1. Validation of the chemical-genetic interactions.** We selected 15 overexpression strains and measured the minimum inhibitory concentration (MIC) of the corresponding AMPs for these strains. In total *n* = 83 MIC measurements were performed. **a,** Out of the 29 resistance cases, 23 (79.31% true positive) were confirmed by the MIC measurements, while a single case showed an opposite effect. **b,** In the case of sensitivity, out of 54 cases, 47 (87.04% true positive) showed the expected MIC change. For details see Supplementary Table 3.

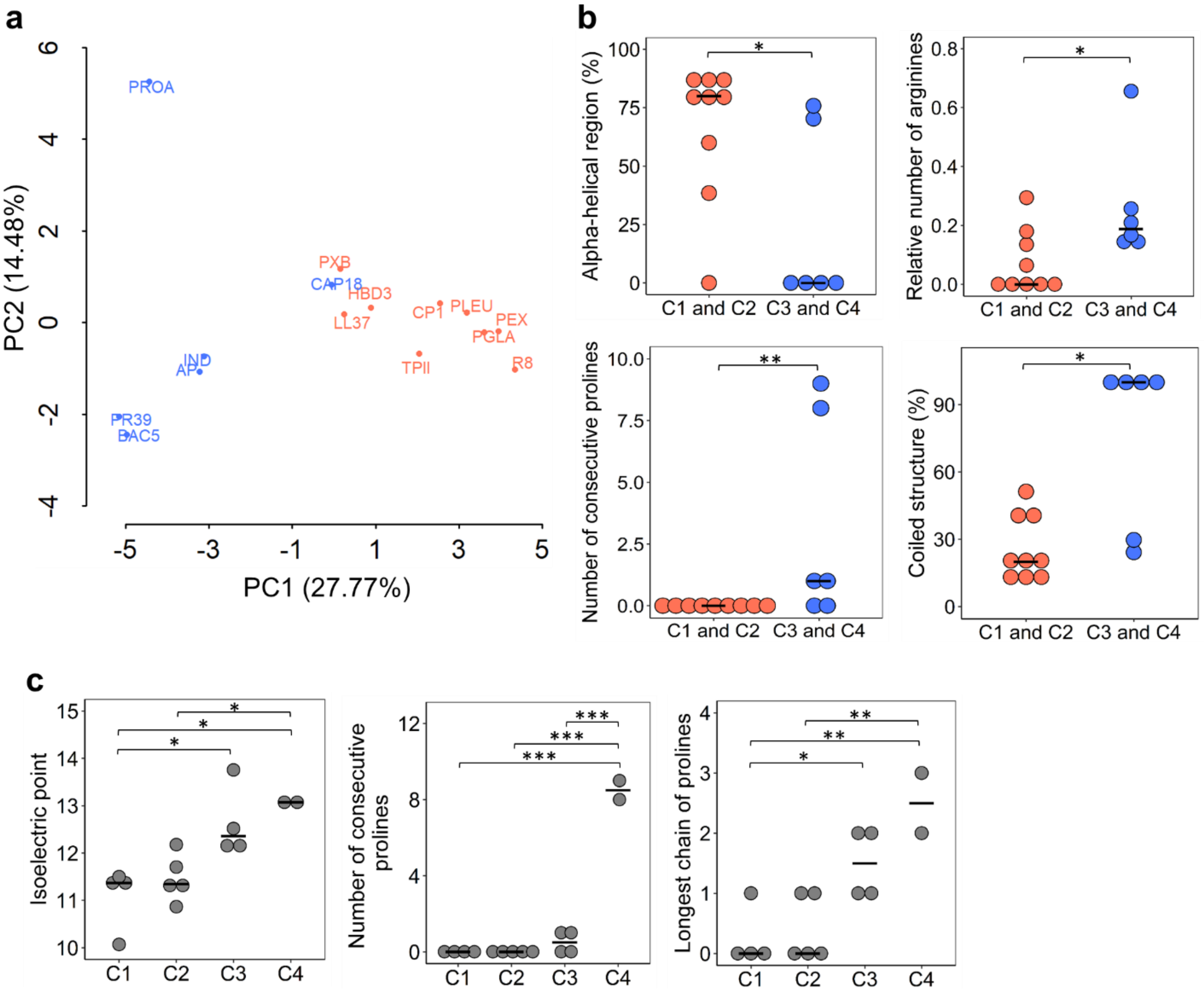


**Supplementary Figure 2. Physicochemical properties of AMPs that differentiate the clusters. a,** Principle component analysis (PCA) of the physicochemical properties differentiated AMPs in C1-C2 clusters (orange) from AMPs in C3-C4 clusters (blue) ($P$ = 4.128*10$^{-5}$, logistic regression, $n$ = 15 AMPs). **b,** Four important physicochemical properties that differentiate membrane-targeting AMPs (orange, C1 and C2) from intracellular-targeting AMPs (blue, C3 and C4). AMPs from C1 and C2 clusters have more alpha-helical regions in their sequence compared to those from C3 and C4 clusters. AMPs from C3 and C4 clusters can be distinguished from those in clusters C1 and C2 based on their higher proline and arginine content. Significant differences from two-sided Mann–Whitney U test, * $P$ = 0.0264, $P$ = 0.0306 and $P$ = 0.0147 for alpha-helical region (%), relative number of arginines and coiled structure (%), respectively. ** $P$ = 0.008 for number of consecutive prolines ($n$ = 9 for C1, C2 and n = 6 for C3, C4). **c,** Physicochemical properties that distinguish the clusters when AMPs in the four clusters are analysed separately (p<0.05 ANOVA, Tukey post hoc test). AMPs in clusters C1 and C2 show an especially low isoelectric point (significant differences: * *P* = 0.017, *P* = 0.014 and *P* = 0.04 for C1 vs C3, C1 vs C4 and C2 vs C4, respectively). AMPs from C4 cluster have an especially high number of consecutive prolines in their amino acid sequences compared to all other clusters (including cluster C3) (significant differences: *** $P = 3.2*10^{-11}$, $P = 9.8*10^{-12}$ and $P = 1.1*10^{-10}$ for C1 vs C4, C2 vs C4 and C3 vs C4 respectively. AMPs from C1 and C2 can also be differentiated based on longest chain of prolines in their sequences (significant differences: * *P* = 0.039 for C1 vs C3. ** *P* = 0.0034 and *P* = 0.0044 for C1 vs C4 and C2 vs C4, respectively, *n* = 4, 5, 4 and 2 for AMPs from C1, C2, C3 and C4 clusters, respectively. Central horizontal bars represent median values. Data is provided in Supplementary Table 5.

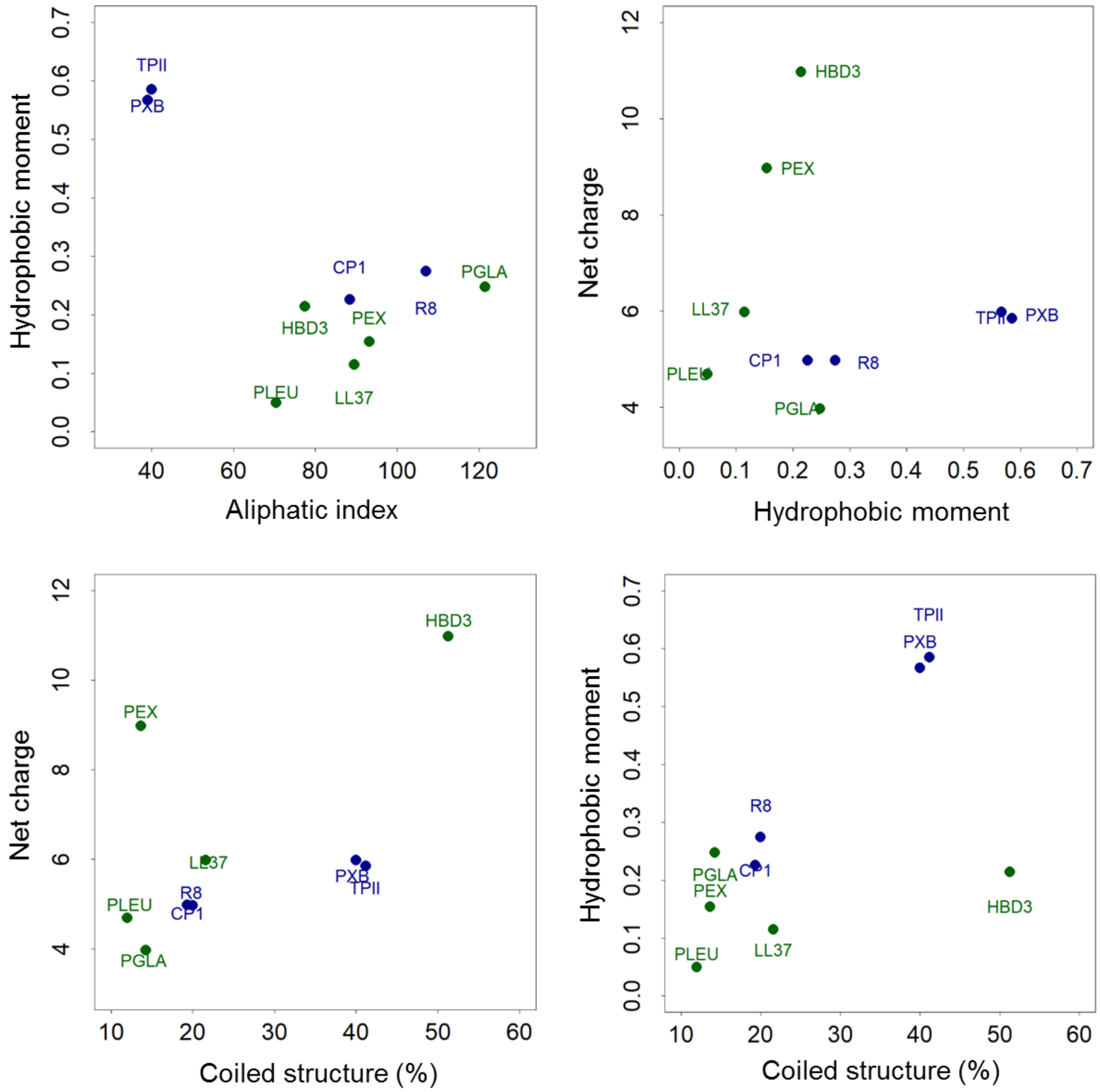


**Supplementary Figure 3. Physicochemical properties of AMPs that jointly separate C1 (blue) and C2 cluster peptides (green).** Although AMPs from both C1 and C2 clusters are known to target the bacterial cell membrane by creating pores in it, they group into different clusters on the basis of their chemical-genetic interactions (see main text and figures). Reassuringly, some physicochemical properties, when considered together, separate AMPs belonging to these two clusters ($P$ = 0.0186 from two-sided logistic regression for all four combinations). $N$ = 4 and 5 for AMPs from C1 and C2 clusters, respectively. Data is provided in Supplementary Table 5.

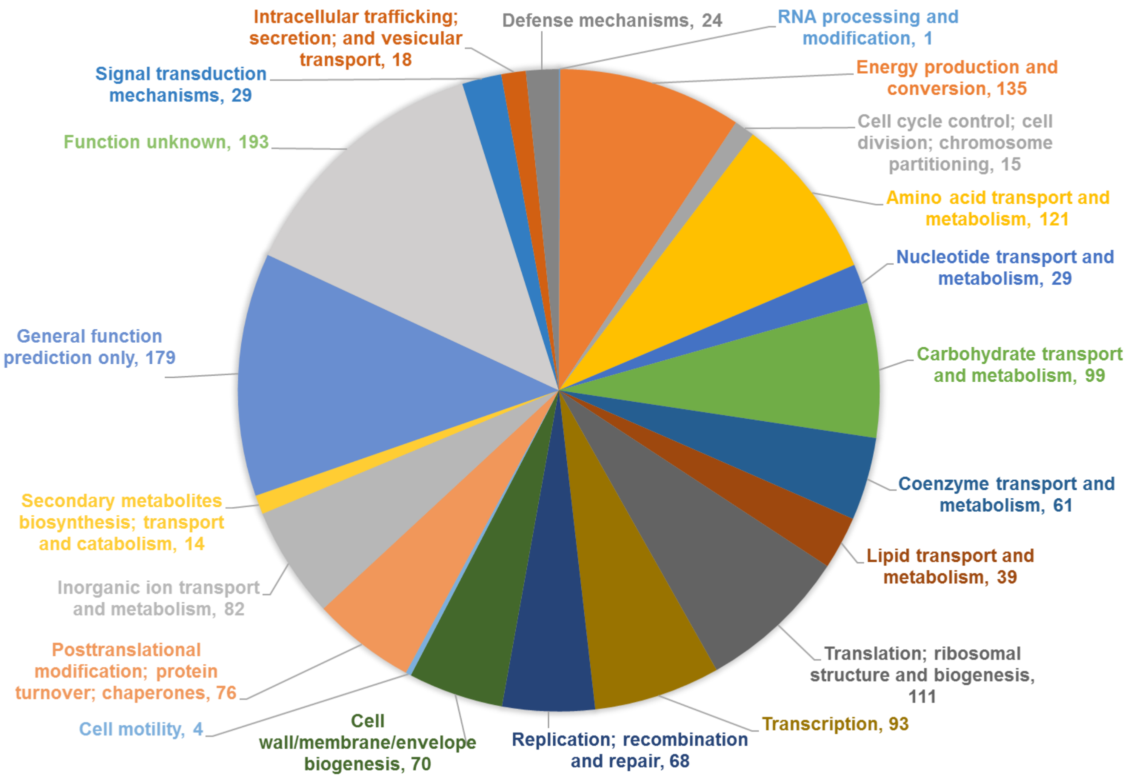


**Supplementary Figure 4.** Distribution of Clusters of Orthologous Groups (COG) categories among the genes that show resistance to at least one AMP in the chemical-genetic screen. Value with each COG category shows total number of genes that belong to corresponding COG category. Data is provided in Supplementary Table 6.

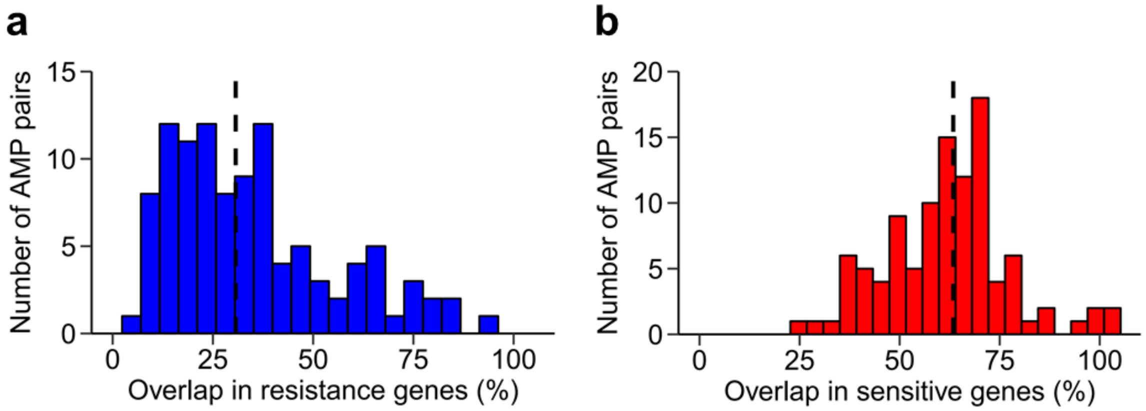


**Supplementary Figure 5. Resistance-conferring genes overlap only to a limited extent between AMPs.** Distributions of overlaps in the **(a)** resistant (blue) and **(b)** sensitive (red) chemical-genetic interactions for all possible AMP pairs (*n* = 105 AMP pairs). Dashed line represents median value. Data is provided in Supplementary Table 6.

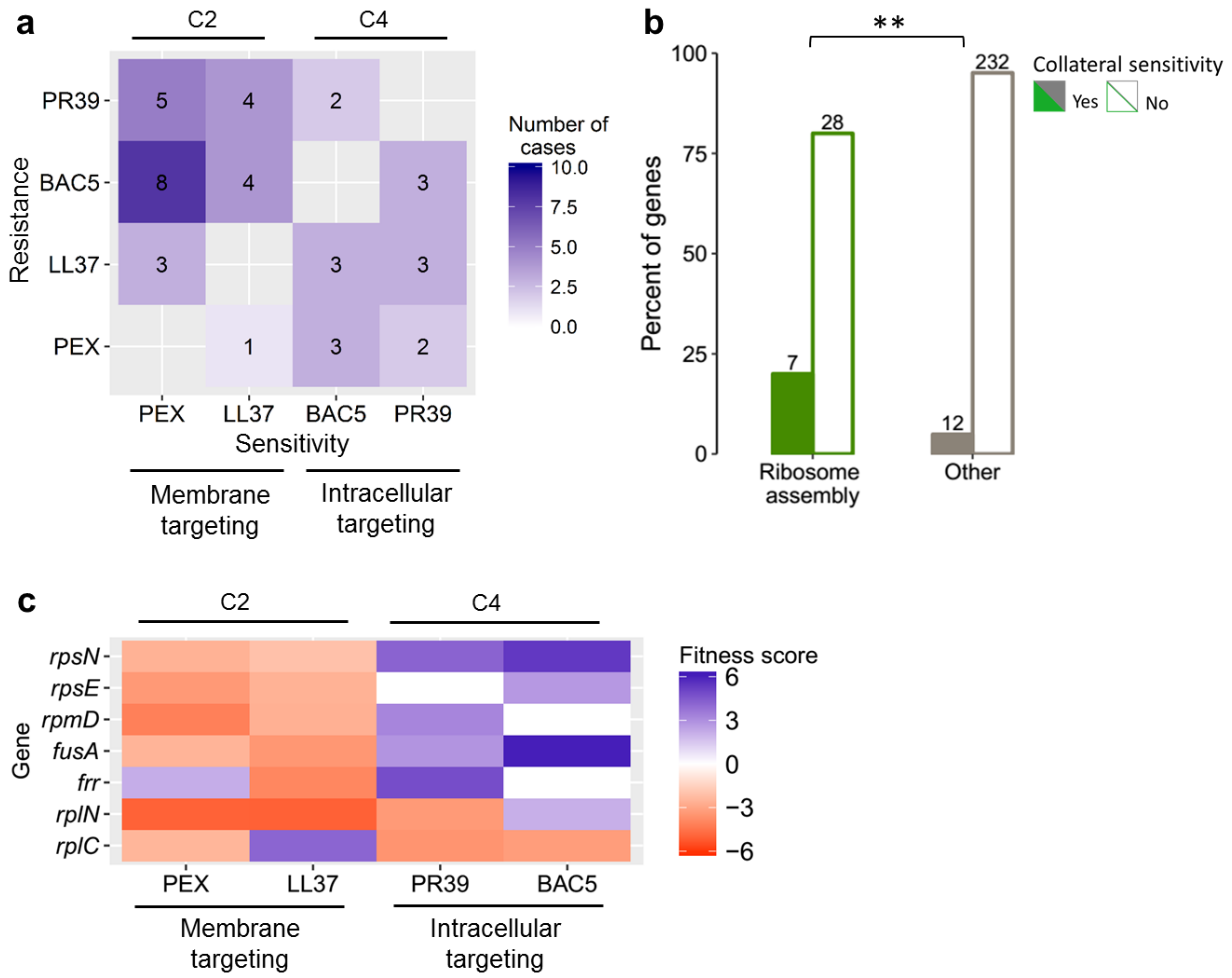


**Supplementary Figure 6. Hypomorphic alleles induce collateral sensitivity between intracellular-targeting and membrane-targeting AMPs. a,** Heatmap shows collateral sensitivity interaction between AMPs. Values in squares represent number of genes that show collateral sensitivity interactions. **b,** Genes showing collateral sensitivity interactions were significantly enriched in functions related to ribosome assembly (significant difference: ** *P* = 0.004 from two-sided Fisher's exact test, *n* = 35 and 244 for ribosome assembly related and other genes, respectively). **c,** Heatmap depicting fitness scores of the genes enriched in ribosome assembly function. Blue and red colors represent resistance and sensitivity interactions, respectively. Data is provided in Supplementary Table 8.

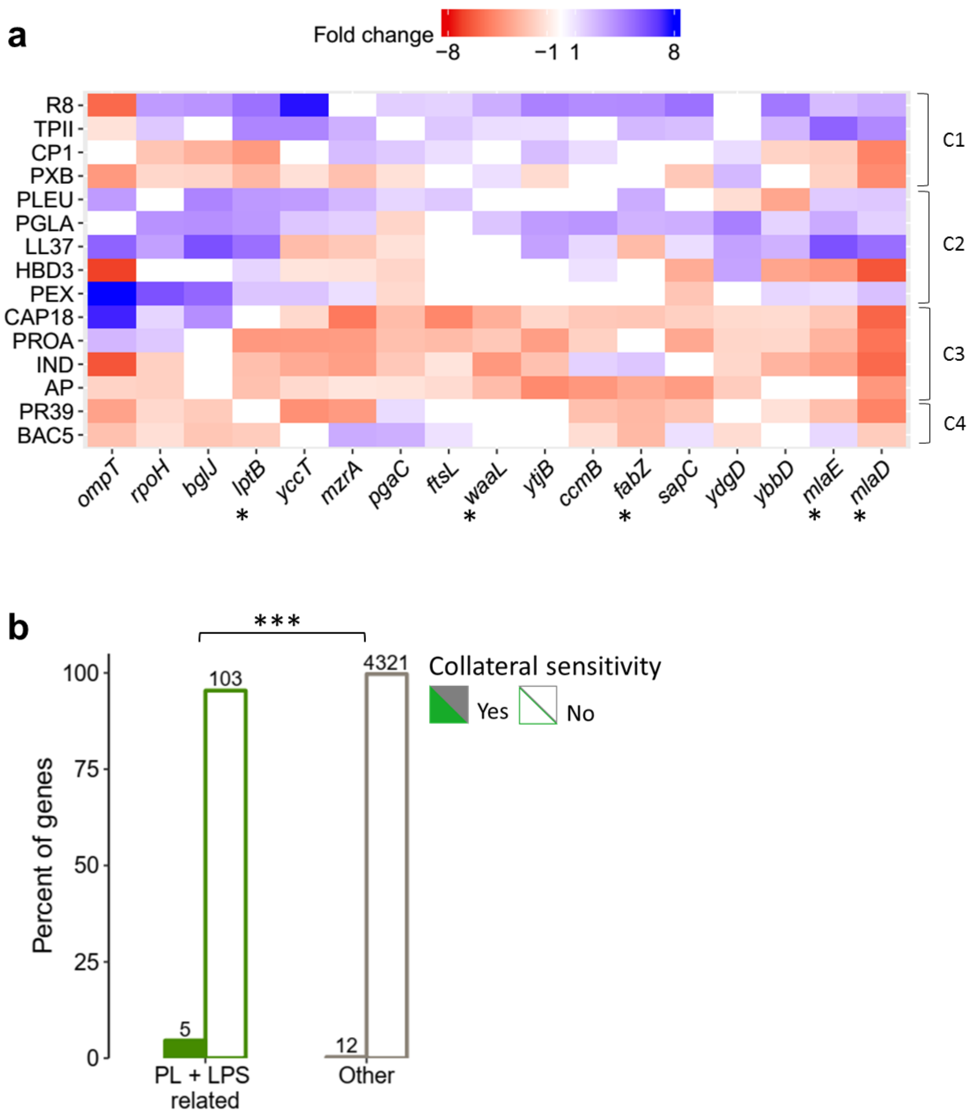


**Supplementary Figure 7. Phospholipid (PL) and lipopolysaccharide (LPS)-related genes frequently show collateral sensitivity interactions. a,** Heatmap presents the chemical-genetic profiles of genes showing resistance (blue) to at least 4 membrane-targeting AMPs and, simultaneously, sensitivity (red) to at least 4 intracellular-targeting AMPs, upon overexpression. Genes marked with asterisk have functions related to PL and LPS composition of the bacterial membranes. **b,** Overexpressed genes showing resistance to at least 4 membrane-targeting AMPs while at the same time sensitivity to at least 4 intracellular-targeting AMPs were significantly enriched in functions related to PL and LPS composition of the bacterial membranes (significant difference: *** $P = 3.8*10^{-5}$ from two-sided Fisher's exact test, $n$ = 108 and 4,333 for PL+LPS related and other genes, respectively). Data is presented in Supplementary Table 6.

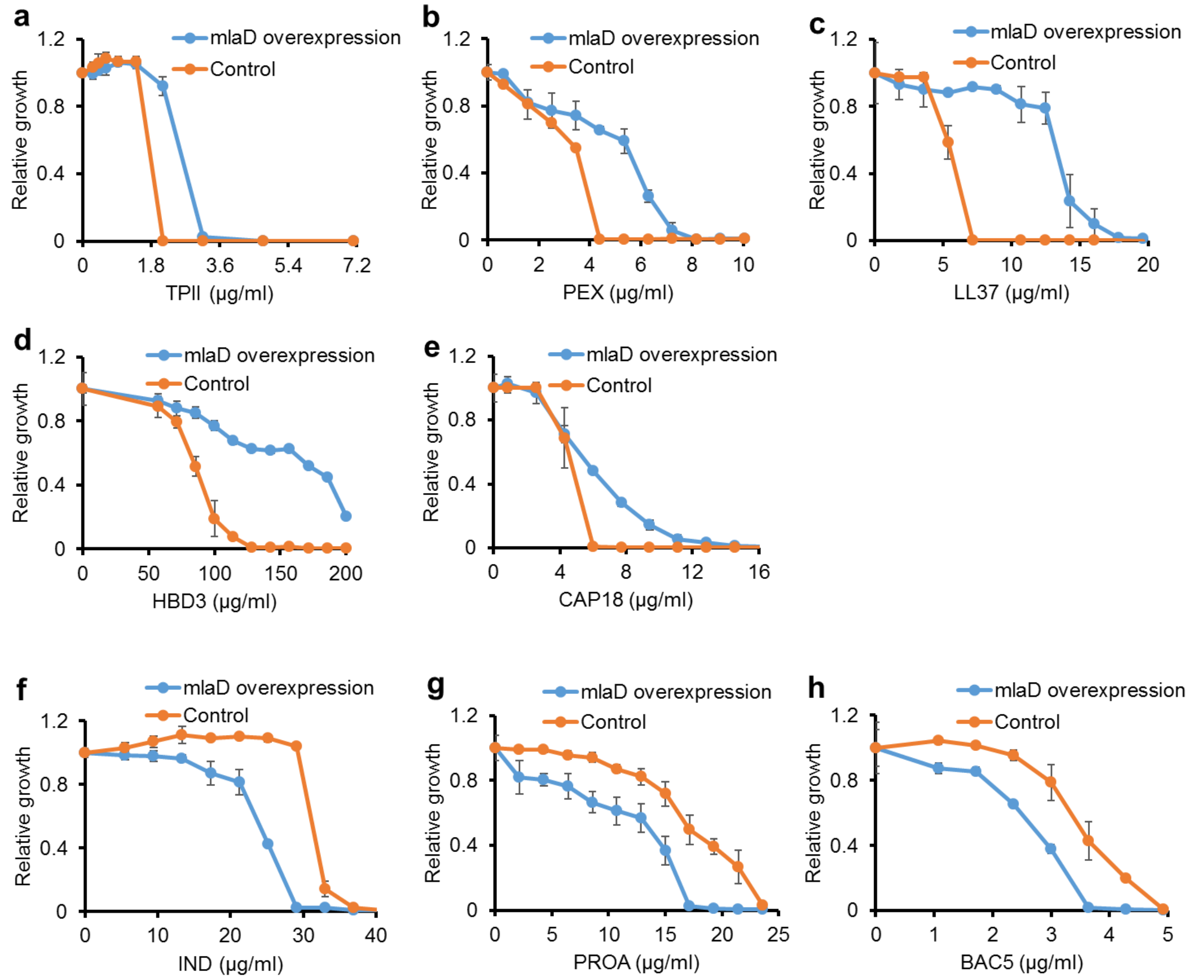


**Supplementary Figure 8. Effect of *mlaD* overexpression on bacterial susceptibility to membrane- and intracellular-targeting AMPs.** Overexpression of *mlaD* causes resistance to membrane-targeting **(a-e)** and sensitivity to intracellular-targeting AMPs **(f-h)**. Bacterial susceptibility to AMPs was tested by measuring MICs (see Methods). Growth is shown relative to growth in the absence of the given AMP (y-axis). The blue line represents the *mlaD* overexpression strain and the orange line represents the control strain (wild-type *E. coli* BW25113 containing the empty pCA24N vector). Error bars indicate standard errors based on three biological replicates.

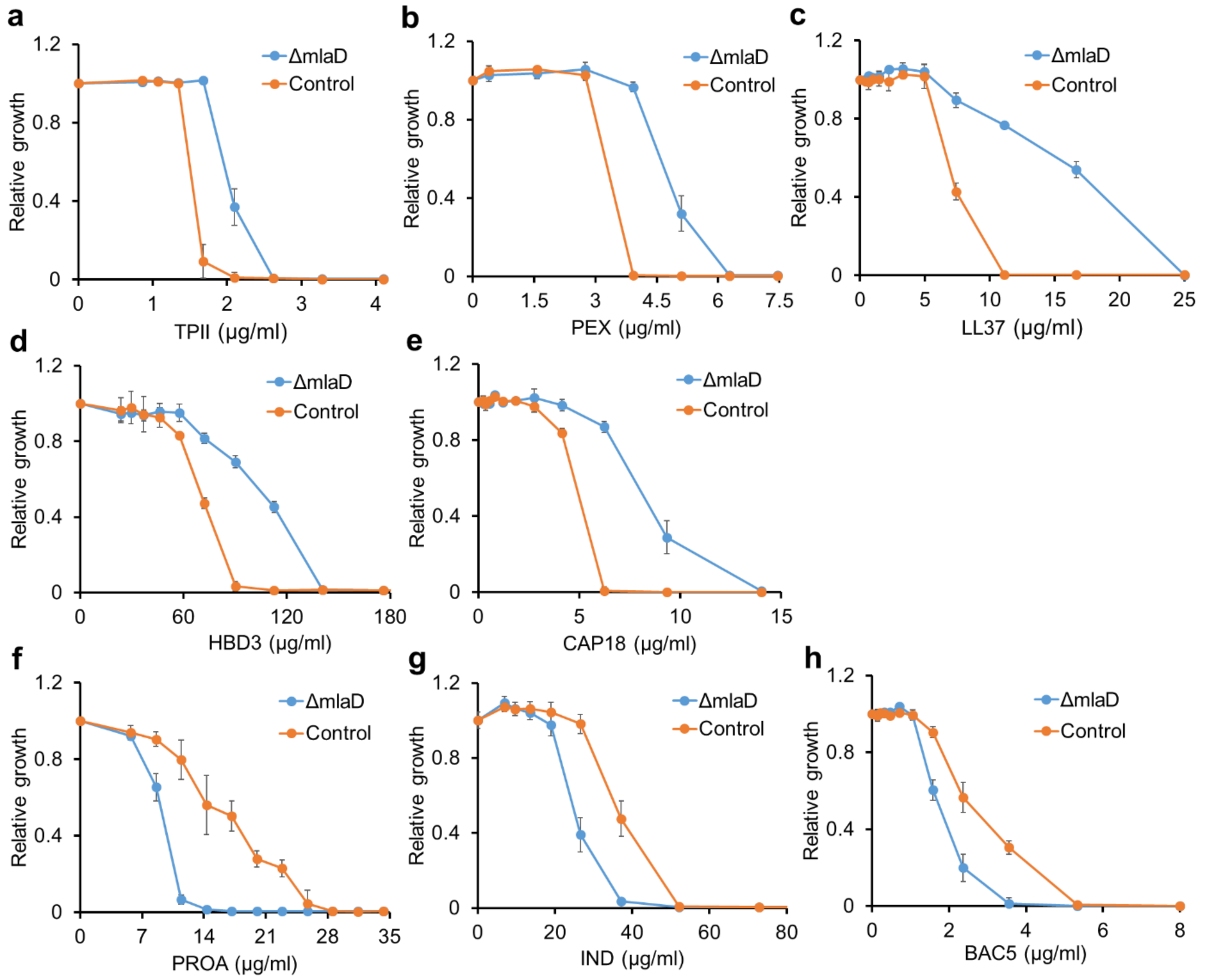


**Supplementary Figure 9. Antimicrobial susceptibility of *mlaD* knockout mutant to membrane- and intracellular-targeting AMPs.** *mlaD* knockout mutant shows resistance to membrane-targeting **(a-e)** and sensitivity to intracellular-targeting AMPs **(f-h)**. The bacterial susceptibility of the mutant to AMPs was tested by MIC determination. Growth is shown relative to growth in the absence of the corresponding AMP. The blue line represents the *mlaD* knockout strain and the orange line represents the control strain (wild-type *E. coli* BW25113). Error bars indicate the standard errors based on three biological replicates.

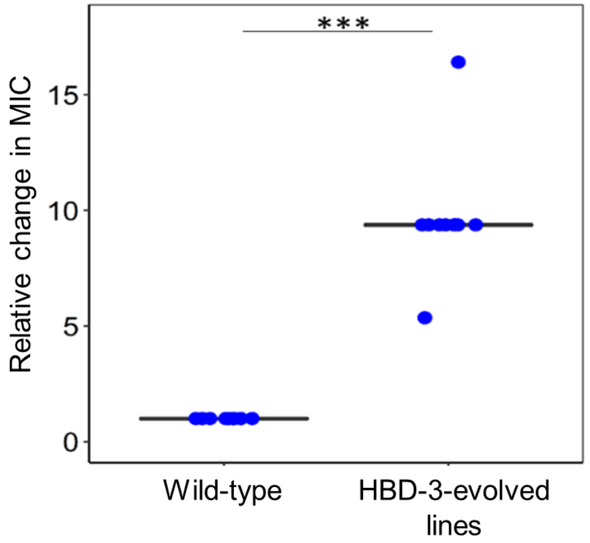


**Supplementary Figure 10. Susceptibility of HBD-3-evolved lines towards HBD-3 AMP.** HBD-3-evolved lines show a significant increase in MIC to HBD-3 as compared to the wild-type ancestral strain (significant difference: *** $P = 3.318*10^{-05}$ from two-sided Mann–Whitney U test, $n = 10$ biological replicates for wild-type ancestral strain and $n = 10$ for HBD-3-lines evolved independently).

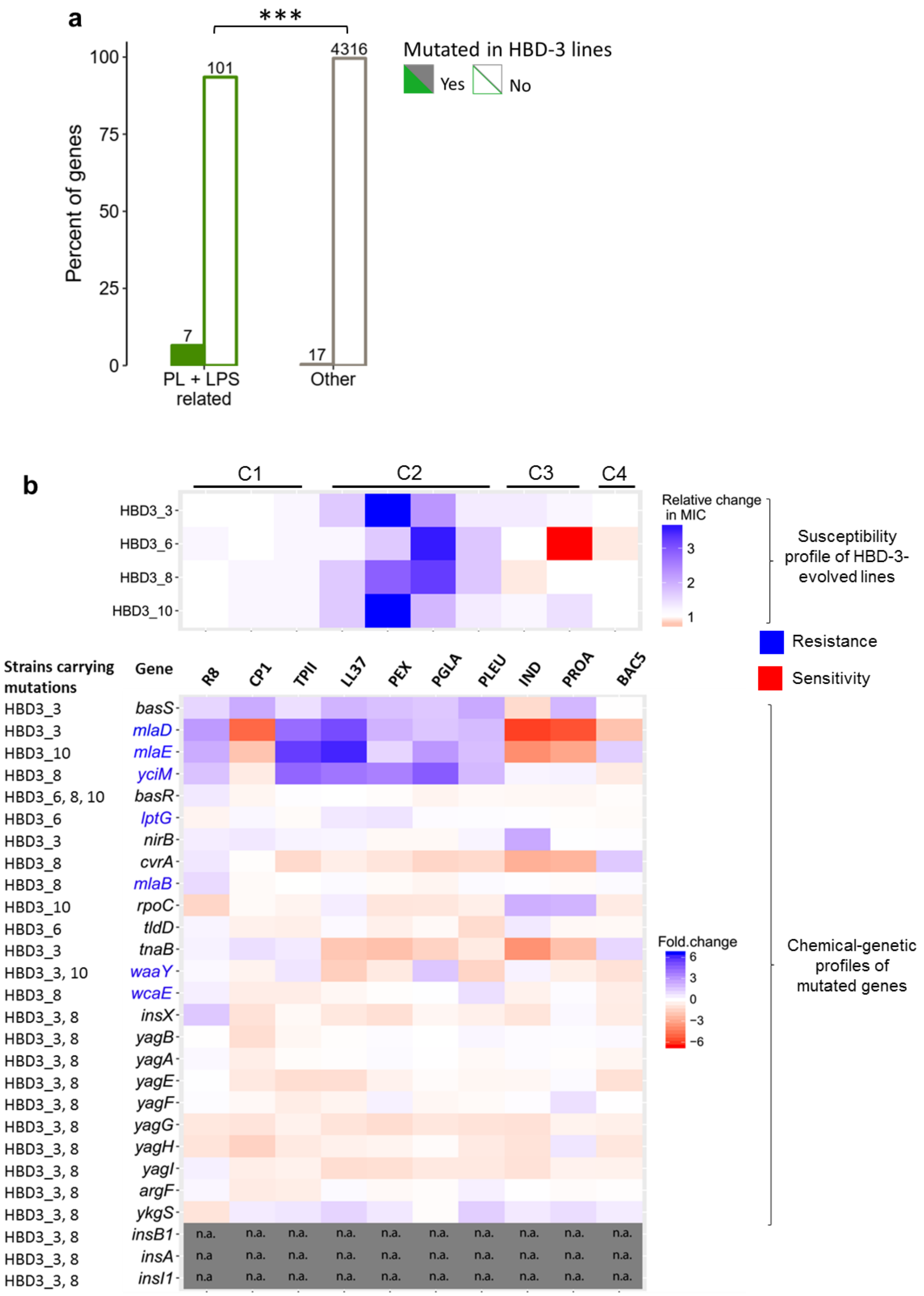
a
***
Percent of genes
100
75
50
25
0
101
4316
7
17
PL + LPS related
Other
Mutated in HBD-3 lines
Yes
No
b
C1
C2
C3
C4
HBD3_3
HBD3_6
HBD3_8
HBD3_10
Relative change in MIC
3
2
1
Susceptibility profile of HBD-3-evolved lines
Resistance
Sensitivity
Strains carrying mutations
Gene
R8
CP1
TPII
LL37
PEX
PGLA
PLEU
IND
PROA
BAC5
HBD3_3 basS
HBD3_3 mlaD
HBD3_10 mlaE
HBD3_8 yciM
HBD3_6, 8, 10 basR
HBD3_6 lptG
HBD3_3 nirB
HBD3_8 cvrA
HBD3_8 mlaB
HBD3_10 rpoC
HBD3_6 tldD
HBD3_3 tnaB
HBD3_3, 10 waaY
HBD3_8 wcaE
HBD3_3, 8 insX
HBD3_3, 8 yagB
HBD3_3, 8 yagA
HBD3_3, 8 yagE
HBD3_3, 8 yagF
HBD3_3, 8 yagG
HBD3_3, 8 yagH
HBD3_3, 8 yagI
HBD3_3, 8 argF
HBD3_3, 8 ykgS
HBD3_3, 8 insB1
HBD3_3, 8 insA
HBD3_3, 8 insI1
n.a.
Fold.change
6
3
0
−3
−6
Chemical-genetic profiles of mutated genes

**Supplementary Figure 11. Chemical-genetic profiles of genes mutated in the HBD-3-evolved lines show similarity to the observed cross-resistance patterns of the evolved lines. a,** Phospholipid and LPS related genes are overrepresented among genes mutated in the HBD-3-evolved lines (significant difference: *** $P = 1*10^{-6}$ from two-sided Fisher's exact test, $n$ = 108 and 4,333 for PL+LPS related genes and others, respectively). **b,** Heatmaps show antimicrobial susceptibility of HBD-3-evolved lines to different AMPs (top heatmap). HBD-3-evolved lines show high cross-resistance to AMPs of C2 cluster (adapted from Figure 6A). Color scale shows the relative change in MIC. Cross-resistance pattern of HBD-3-evolved lines showed similarity with chemical-genetic profiles of the genes (bottom heatmap) mutated in these evolved lines. Phospholipid (PL) and lipopolysaccharide (LPS)-related genes are highlighted with blue color. Data is presented in Supplementary Table 6.

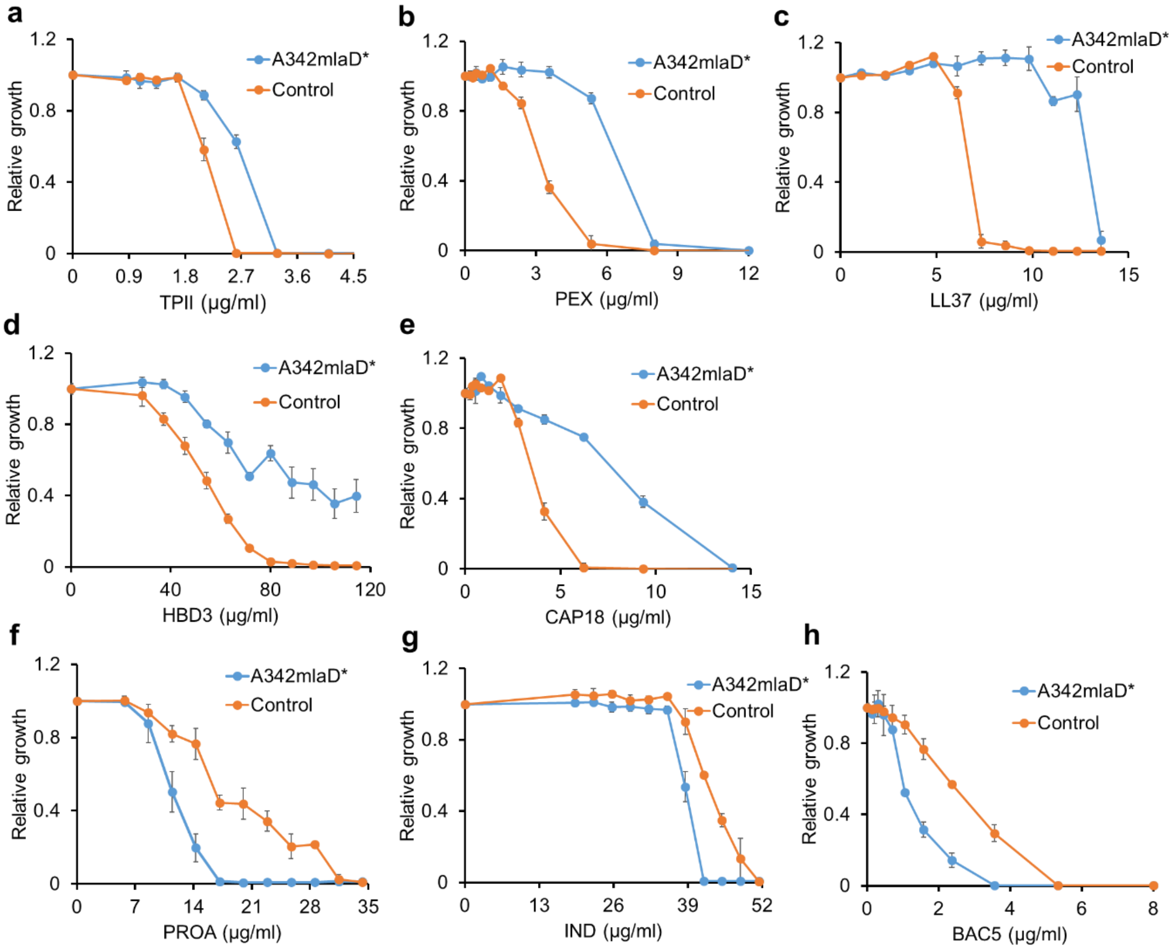


**Supplementary Figure 12. Antimicrobial susceptibility of *A342mlaD** mutant to membrane- and intracellular-targeting AMPs.** The A342mlaD* mutant shows resistance to membrane-targeting **(a-e)** and sensitivity to intracellular-targeting AMPs **(f-h)**. Effect of the mutation on bacterial susceptibility to AMPs was tested by MIC measurements (see Methods). Growth is shown relative to growth in the absence of AMP. Error bars indicate standard errors based on three biological replicates. The blue line represents the *A342mlaD** mutant strain and the orange line represents the control strain (wild-type *E. coli* BW25113).

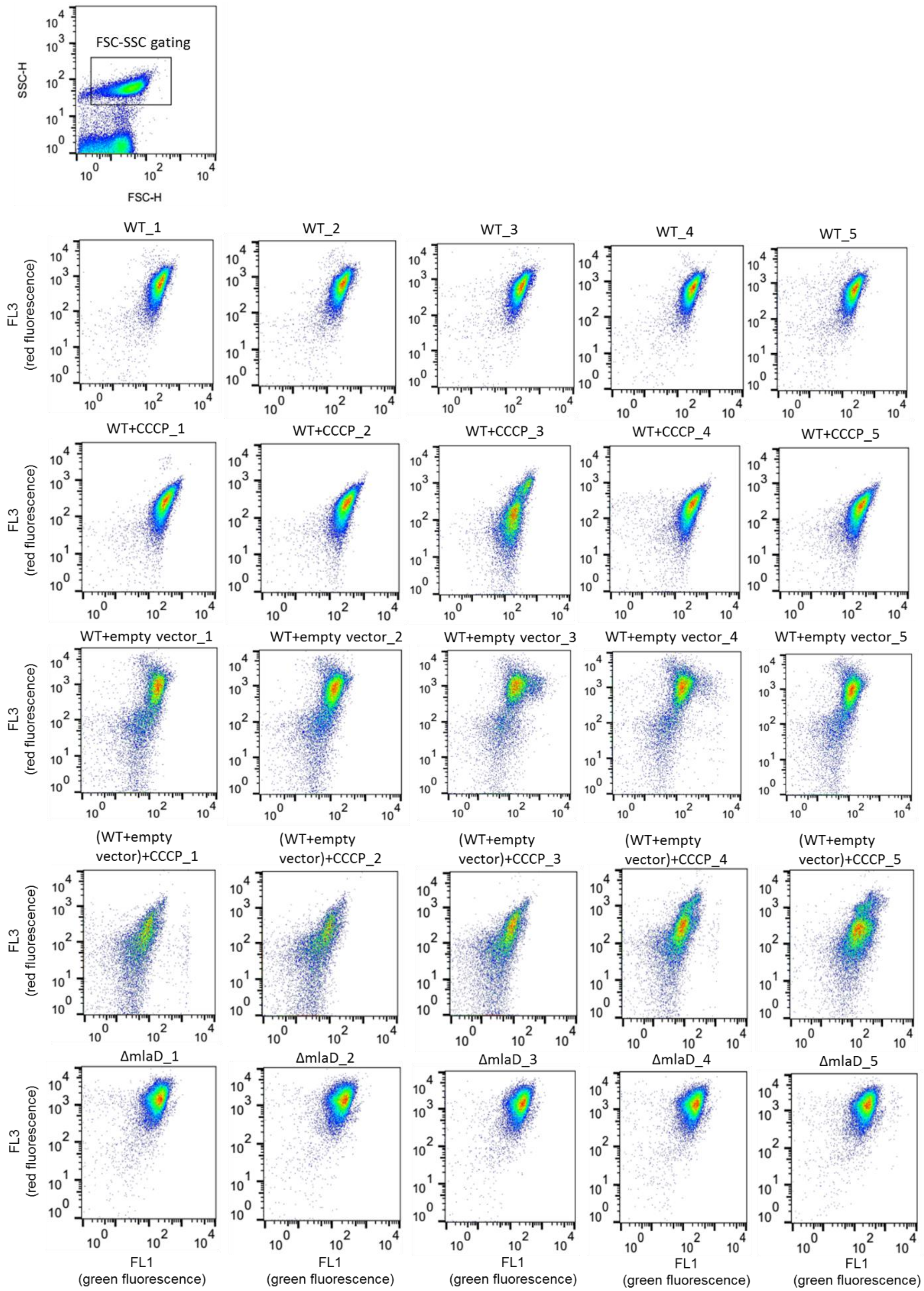
FSC-SSC gating
SSC-H
FSC-H
WT_1
WT_2
WT_3
WT_4
WT_5
WT+CCCP_1
WT+CCCP_2
WT+CCCP_3
WT+CCCP_4
WT+CCCP_5
WT+empty vector_1
WT+empty vector_2
WT+empty vector_3
WT+empty vector_4
WT+empty vector_5
(WT+empty vector)+CCCP_1
(WT+empty vector)+CCCP_2
(WT+empty vector)+CCCP_3
(WT+empty vector)+CCCP_4
(WT+empty vector)+CCCP_5
ΔmlaD_1
ΔmlaD_2
ΔmlaD_3
ΔmlaD_4
ΔmlaD_5
FL3 (red fluorescence)
FL1 (green fluorescence)

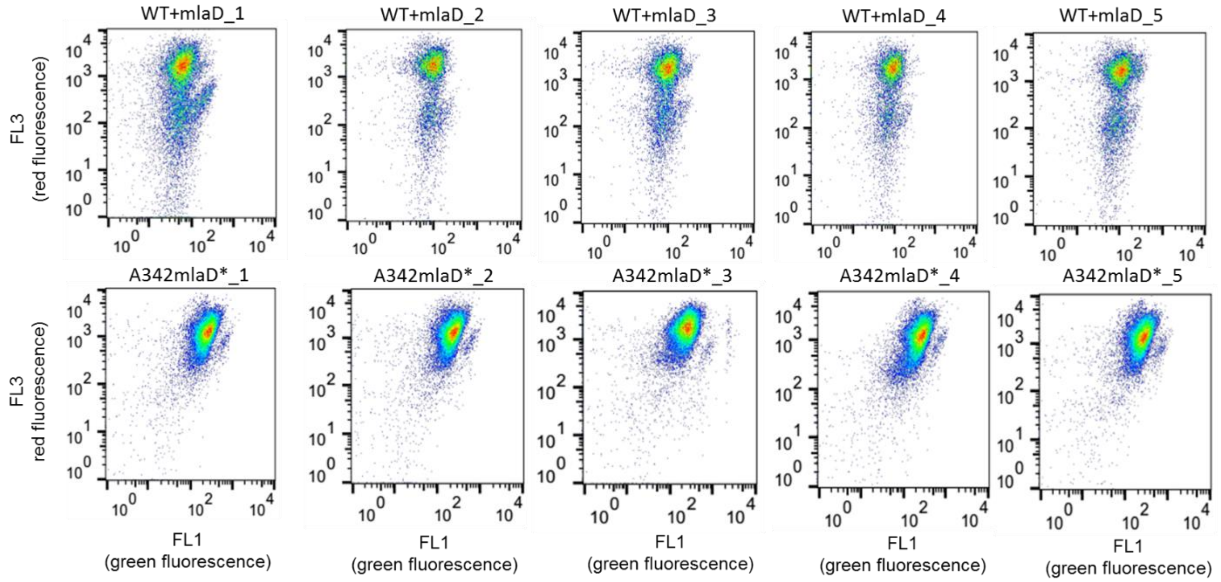


**Supplementary Figure 13**. Membrane potential measurement. Red (FL3-H) vs green (FL1-H) fluorescence scatter plots showing cells treated with the $DiOC_2(3)$ dye (membrane potential indicator, see Methods). For each strain we had five biological replicates. Top figure showed gating strategy where cells were gated using forward and side scatter properties. Data is present in Supplementary Table 12.

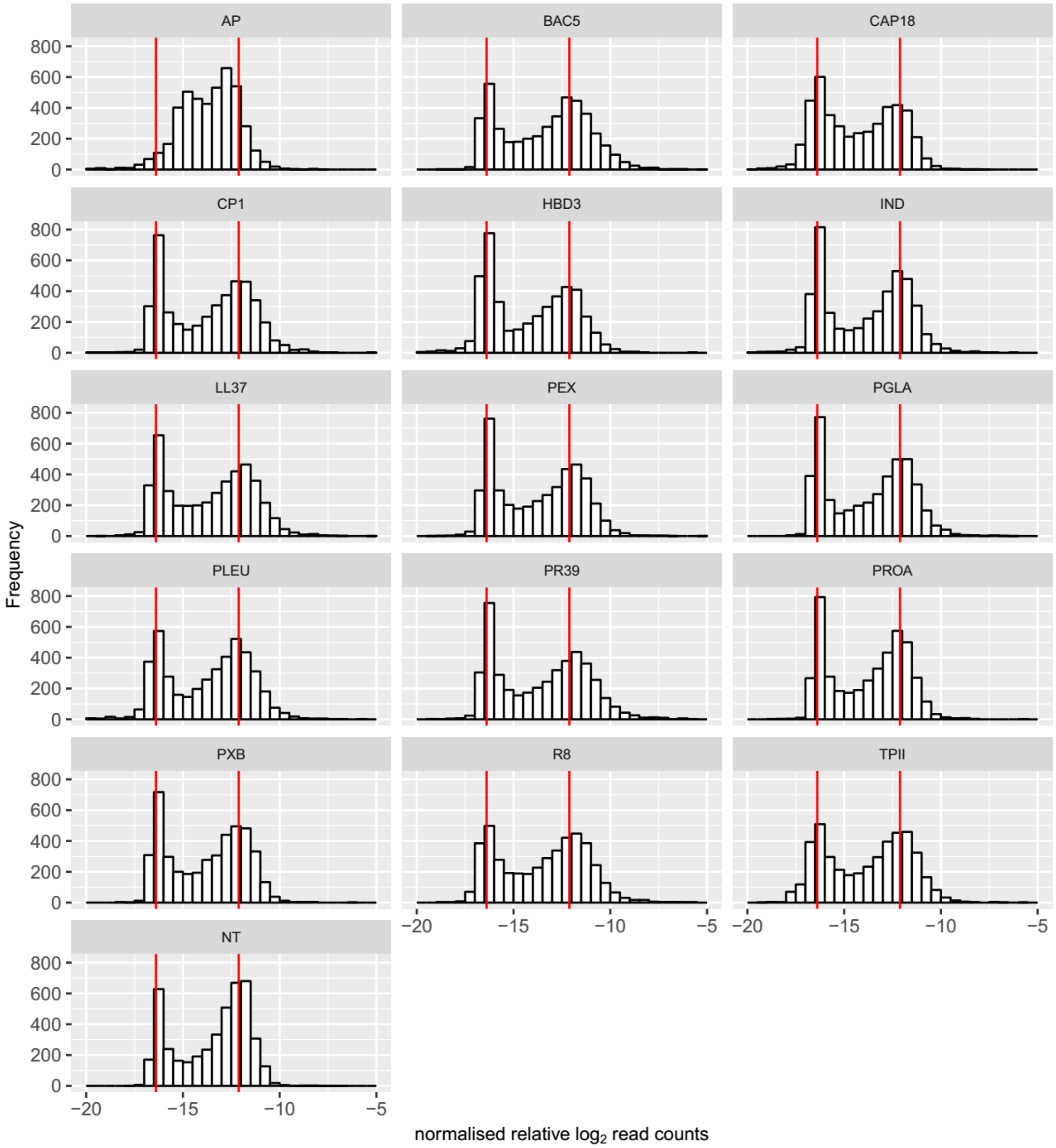


**Supplementary Figure 14**. Distribution of the normalized read counts of plasmids overexpressing *E. coli* ORFs in the absence (NT) and presence of AMPs. Following data processing (see Methods), the transformed relative read counts showed bimodal distributions (peaks of the two modes are marked with red lines). The lower mode of the distribution corresponds to ORFs that were not present in the sample. The upper mode represents those ORFs whose growth was unaffected by overexpression (i.e. no fitness effect).

### Supplementary Tables

**Supplementary Table 1. A comprehensive catalogue of previously reported genes (based on literature mining) that modulate bacterial susceptibility to antimicrobial peptides (AMPs).**

Provided in a separate Excel spreadsheet.

**Supplementary Table 2. Complete dataset of chemical-genetic interactions of ~4400 mutants across 15 different antimicrobial peptides (AMPs).**

Provided in a separate Excel spreadsheet.

**Supplementary Table 3. Relative changes in the minimum inhibitory concentrations (MIC) of the antimicrobial peptides (AMPs) towards 15 overexpression strains**.

Provided in a separate Excel spreadsheet.

**Supplementary Table 4. Collected examples of *E. coli* gene overexpression from the literature as a confirmation of our chemical-genetic screen results.**

| Gene | AMP | Overexpression phenotype | |
|---|---|---|---|
| | | Literature | Chemical-genetic profile |
| *ptrB* | BAC5 | Resistance[1] | Confirmed |
| | PR39 | Resistance[1] | Confirmed |
| *marA* | PXB | Resistance[2] | Confirmed |
| | LL37 | Resistance[2] | X |
| *ompT* | PROA | Resistance[3] | Confirmed |
| | LL37 | Resistance[3] | Confirmed |
| *arnT* | PXB | Resistance[4] | X |
| *nlpE* | PROA | Resistance[5] | Confirmed |
| *eptA* | PXB | Resistance[6] | Confirmed |
| *pmrD* | PXB | Resistance[7] | X |
| *sbmA* | PR39 | Sensitivity[8] | Confirmed |
| *waaY* | CAP18 | Sensitivity[9] | Confirmed |
| *prfA* | AP | Resistance[10] | X |

**Supplementary Table 5. List of the physicochemical properties of the antimicrobial peptides (AMPs) used in this study.**

Provided in a separate Excel spreadsheet.

**Supplementary Table 6. List of resistance- and sensitivity-conferring genes and their collateral sensitivity (CS) interactions.**

Provided in a separate Excel spreadsheet.

**Supplementary Table 7. Gene ontology (GO) enrichment analysis of the genes conferring resistance and sensitivity to antimicrobial peptides (AMPs).**

Provided in a separate Excel spreadsheet.

**Supplementary Table 8. Complete dataset of chemical-genetic interactions of hypomorphic alleles and antimicrobial peptides (AMPs).**

Provided in a separate Excel spreadsheet.

**Supplementary Table 9. List of genes mutated in HBD-3-evolved lines.**

| Strain | Type | Gene | Mutation | Description |
|---|---|---|---|---|
| HBD3_3 | Deletion | *[insI1]–insA* | Δ11,471 bp | Genes affected by the deletion: [insI1], insX, yagB, yagA, yagE, yagF, yagG, yagH, yagI, argF, ykgS, insB1, insA |
| HBD3_8 | Deletion | *[insI1]–insA* | Δ11,471 bp | Genes affected by the deletion: [insI1], insX, yagB, yagA, yagE, yagF, yagG, yagH, yagI, argF, ykgS, insB1, insA |
| HBD3_10 | SNP | *basR* | C→T | Response regulator in two component regulatory system with BasS |
| HBD3_6 | SNP | *basR* | G→T | Response regulator in two component regulatory system with BasS |
| HBD3_8 | SNP | *basR* | C→A | Response regulator in two component regulatory system with BasS |
| HBD3_3 | SNP | *basS* | A→C | Sensory histidine kinase in two component regulatory system with BasR |
| HBD3_8 | SNP | *cvrA* | A→C | Putative cation/proton antiporter |
| HBD3_6 | SNP | *lptG* | G→A | Lipopolysaccharide export ABC permease of the LptBFGC export complex |
| HBD3_8 | SNP | *mlaB* | G→A | ABC transporter maintaining OM lipid asymmetry |

| | | | | |
|---|---|---|---|---|
| HBD3_3 | Deletion | *mlaD* | Δ1 bp | ABC transporter maintaining OM lipid asymmetry |
| HBD3_10 | SNP | *mlaE* | G→T | ABC transporter maintaining OM lipid asymmetry |
| HBD3_3 | SNP | *nirB* | C→A | Nitrite reductase, large subunit, NAD(P)H binding |
| HBD3_10 | SNP | *rpoC* | T→G | RNA polymerase, beta prime subunit |
| HBD3_6 | SNP | *tldD* | A→C | Putative peptidase |
| HBD3_3 | Insertion | *tnaB* | IS5 (–) +4 bp | Tryptophan transporter of low affinity |
| HBD3_10 | SNP | *waaY* | T→G | Lipopolysaccharide core biosynthesis protein |
| HBD3_3 | SNP | *waaY* | T→G | Lipopolysaccharide core biosynthesis protein |
| HBD3_8 | Insertion | *wcaE* | IS5 (+) +4 bp | Putative glycosyl transferase |
| HBD3_8 | SNP | *yciM* | G→A | Envelope integrity maintenance protein |
| HBD3_6 | SNP | *yneO* ← / ← *lsrK* | T→C | Pseudogene, AidA homolog/autoinducer2 (AI2) kinase |

**Supplementary Table 10. Detailed information (based on literature mining) of the antimicrobial peptides (AMPs) used in this study.**

Provided in a separate Excel spreadsheet.

**Supplementary Table 11. Susceptibility profiles of HBD-3-evolved lines across different antimicrobial peptides (AMPs).**

Provided in a separate Excel spreadsheet.

**Supplementary Table 12. Raw dataset of red (FL3-H) and green (FL1-H) fluorescence from membrane potential measurement.**

Provided in a separate Excel spreadsheet.

**References (Supplementary)**


1. Mattiuzzo, M. *et al.* Proteolytic Activity of Escherichia coli Oligopeptidase B Against Proline-Rich Antimicrobial Peptides. *J. Microbiol. Biotechnol.* **24**, 160–167 (2014).
2. Warner, D. M. & Levy, S. B. Different effects of transcriptional regulators MarA, SoxS and Rob on susceptibility of Escherichia coli to cationic antimicrobial peptides (CAMPs): Rob-dependent CAMP induction of the marRAB operon. *Microbiology* **156**, 570–578 (2010).
3. Stumpe, S., Schmid, R., Stephens, D. L., Georgiou, G. & Bakker, E. P. Identification of OmpT as the protease that hydrolyzes the antimicrobial peptide protamine before it enters growing cells of Escherichia coli. *J. Bacteriol.* **180**, 4002–6 (1998).
4. Trent, M. S. *et al.* Accumulation of a Polyisoprene-linked Amino Sugar in Polymyxin-resistant Salmonella typhimurium and Escherichia coli. *J. Biol. Chem.* **276**, 43132–43144 (2001).
5. Identification of Novel Genetic Mechanisms Required for Bacterial Resistance to Antimicrobial Peptides**.** (https://repository.asu.edu/attachments/110499/content/Griffin_asu_0010E_12909.pdf)
6. Olaitan, A. O., Morand, S. & Rolain, J.-M. Mechanisms of polymyxin resistance: acquired and intrinsic resistance in bacteria. *Front. Microbiol.* **5**, 643 (2014).
7. Roland, K. L., Esther, C. R. & Spitznagel, J. K. Isolation and characterization of a gene, pmrD, from Salmonella typhimurium that confers resistance to polymyxin when expressed in multiple copies. *J. Bacteriol.* **176**, 3589–3597 (1994).
8. Pranting, M., Negrea, A., Rhen, M. & Andersson, D. I. Mechanism and Fitness Costs of PR-39 Resistance in Salmonella enterica Serovar Typhimurium LT2. *Antimicrob. Agents Chemother.* **52**, 2734–2741 (2008).
9. Lázár, V. *et al.* Antibiotic-resistant bacteria show widespread collateral sensitivity to antimicrobial peptides. *Nat. Microbiol.* **3**, 718–731 (2018).
10. Matsumoto, K. *et al.* In vivo target exploration of apidaecin based on Acquired Resistance induced by Gene Overexpression (ARGO assay). *Sci. Rep.* **7**, 12136 (2017).
